# Functional Genetic Biomarkers of Alzheimer’s Disease and Gene Expression from Peripheral Blood

**DOI:** 10.1101/2021.01.15.426891

**Authors:** Andrew Ni, Amish Sethi, for the Alzheimer’s Disease Neuroimaging Initiative

**Affiliations:** Pine-Richland High School

**Author notes:** Joint first authors (Andrew Ni, Amish Sethi). Data used in preparation of this article were obtained from the Alzheimer’s Disease Neuroimaging Initiative (ADNI) database (adni.loni.usc.edu). As such, the investigators within the ADNI Contributed to the design and implementation of ADNI and/or provided data but did not participate in analysis or writing of this report. A Complete listing of ADNI investigators can be found at:http://adni.loni.usc.edu/wp-Content/uploads/how_to_apply/ADNI_Acknowledgement_List.pdf.

**Keywords:** Peripheral Blood, Random Forest, Unsupervised Clustering, Causal Analysis, GSEA

## Abstract

Detecting Alzheimer’s Disease (AD) at the earliest possible stage is key in advancing AD prevention and treatment but is challenged by normal aging processes in addition to other confounding neurodegenerative diseases. Recent genome-wide association studies (GWAS) have identified associated alleles, but it has been difficult to transition from non-coding genetic variants to underlying mechanisms of AD. Here, we sought to reveal functional genetic variants and diagnostic biomarkers underlying AD using machine learning techniques. We first developed a Random Forest (RF) classifier using microarray gene expression data sampled from the peripheral blood of 744 participants in the Alzheimer’s Disease Neuroimaging Initiative (ADNI) cohort. After initial feature selection, 5-fold cross-validation of the 100-gene RF classifier achieved an accuracy of 99.04%. The high accuracy of the RF classifier supports the possibility of a powerful and minimally invasive tool for screening of AD. Next, unsupervised clustering was used to validate and identify relationships among differentially expressed genes (DEGs) the RF selected revealing 3 distinct AD clusters. Results suggest downregulation of global sulfatase and oxidoreductase activities in AD through mutations in SUMF1 and SMOX respectively. Then, we used Greedy Fast Causal Inference (GFCI) to find potential causes of AD within DEGs. In the causal graph, HLA-DPB1 and CYP4A11 emerge as hub genes, furthering the discussion of the immune system’s role in AD. Finally, we used Gene Set Enrichment Analysis (GSEA) to determine the biological pathways and processes underlying the DEGs that were highly correlated with AD. Cell activation in the immune system, glycosaminoglycan (GAG) binding, vascular dysfunction, oxidative stress, and the neuronal apoptotic process were revealed to be significantly enriched in AD. This study further advances the possibility of low-cost and noninvasive genetic screening for AD while also providing potential gene targets for further experimentation.

## 1. INTRODUCTION

AD is a neurodegenerative disorder affecting an estimated 6% of all people worldwide that are aged 65 or older [1]. It kills approximately 480,000 people each year [2] and leads to a burden of about $172 billion annually to the health-care system in the US alone [3]. Thus, early detection and treatment has become a priority for our healthcare system. Due to its progressive nature, hallmarks of AD may appear up to 20-30 years before clinical onset [51, 52]. This makes preclinical diagnosis of AD a possibility since pathological symptoms appear years before clinical manifestation [4]. These pathological features include the accumulation of amyloid beta plaques, tau proteins, and neurofibrillary tangles [5, 6]. Despite this possibility of early detection, it has been difficult to develop a system for preclinical diagnosis due to confounding by normal aging processes and other neurodegenerative diseases, in addition to the biological complexity of AD.

Genetics may be the path to a reliable system for early detection. Genetics play a large role in AD and explain an estimated 70% of the risk of development [8]. Currently, Genome Wide Association Studies (GWAS) are one of the most common methods of finding candidate genes of AD. These studies have revealed several genes potentially associated with AD. However, other than the four exceptions of APP, PS-1, PS-2, and APOE, GWAS have failed to produce reliable candidate genes for AD, despite more than thousands of genes being recommended as potential risk factors [9, 48]. This is because GWAS have several limitations. First, GWAS only reveal genes that are associated with a certain phenotype and fail to address which genes functionally cause AD [10]. Second, GWAS fail to account for epistasis, which is the phenomenon by which a complex phenotype is caused by more than one gene interacting. Therefore, few GWAS studies have focused on how genes interact and how this interaction contributes to the progression of AD [47]. For these reasons, GWAS have largely failed in detecting functionally relevant genes of AD.

Recently, studies have turned towards gene expression to identify new genetic biomarkers and create diagnostic tools for AD [53]. These studies utilize gene expression values from the brain tissue in AD subjects. Overall, this means that the analysis was largely based on samples from biopsies or autopsies [54]. This is undesirable because these lab-based samples and analyses are difficult to extrapolate to a clinical setting. In addition, protein modification and RNA degradation postmortem makes it difficult to interpret gene expression from the brain. Instead, expression profiling using peripheral blood yields certain benefits [14].

Peripheral blood mononuclear cells (PBMCs) is easily obtainable and relatively inexpensive, allowing for genetic screening of peripheral blood to be part of the clinical process to assess the risk of AD in living patients. Additionally, changes in the brain are represented through mechanisms that are also expressed in blood. For example, amyloid precursor protein expression, deregulated cytokine secretion, and oxidative damage to DNA and RNA are all shared by AD brain tissue and peripheral blood [18]. Furthermore, more than 80% of the genes expressed in body tissue types, such as brain tissue, are also found in peripheral blood mononuclear cells (PBMCs) [16]. This supports the utilization of peripheral blood to understand molecular mechanisms behind AD. Indeed, in the past, peripheral blood has already been implemented for gene expression analysis for AD with promising results [15, 17].

Therefore, peripheral blood gene expression could be used to advance early detection of AD, both by yielding a diagnostic tool and easily identifiable biomarkers. Here, we revealed functional genetic variants and diagnostic biomarkers underlying AD. In particular, we sought to construct a blood gene expression classifier of AD that could distinguish people with AD from normal control subjects and reveal diagnostic features.

To do this, we first developed a Random Forest (RF) classifier using blood microarray gene expression data from Alzheimer’s Disease Neuroimaging Initiative (ADNI) cohort. Then, unsupervised clustering using Uniform Manifold Approximation and Projection (UMAP) and Hierarchical Density-Based Spatial-Clustering of Applications with Noise (HDBSCAN) were used to identify AD subclusters and reveal relationships among differentially expressed genes (DEGs) the RF selected. After that, causal analysis through the Greedy Fast Causal Inference (GFCI) search algorithm was done to find potential causes of AD within DEGs. Finally, we used Gene Set Enrichment Analysis (GSEA) to determine the biological pathways and processes underlying the DEGs. This study further advances understanding of molecular mechanisms underlying AD and provides potential gene targets for further experimentation.

## 2. DATA

### 2.1 Alzheimer’s Disease Neuroimaging Initiative

Data used for this study was acquired from the Alzheimer’s Disease Neuroimaging Initiative (ADNI) database (adni.loni.usc.edu). Throughout its lifetime, ADNI has enlisted more than 1,500 adults, ages 55 to 90.

For this study, we further scaled and centered the gene expression values for the 744 available individuals. We elected not to collapse the probe sets into individual genes as some probes may not have associated gene annotations, some probes (promiscuous probes) may recognize multiple target sequences, and some genes may have multiple probes. We calculate gene expression by taking the median of the probes for a certain gene.

### 2.3 Oversampling

Training and testing of machine learning algorithms can be heavily impacted by the problem of an imbalanced dataset. This balancing issue arises when the dataset has a significant difference among the size of each class. This imbalance heavily skews the decision function and results in poor training and testing outcomes [20]. With a greater imbalanced ratio, the decision function of the model favors the larger class during classification.

In our case, the ADNI data is clearly imbalanced. We used the diagnoses as the four classes in our dataset. The distribution of classes is: 260 CN, 215 EMCI, 226 LMCI, and 43 AD individuals. AD is underrepresented compared to other groups with only 43 samples.

Synthetic Minority Oversampling Technique (SMOTE) was proposed to address this imbalanced dataset issue by generating new “synthetic” samples of underrepresented classes through interpolation [21]. In this method, a minority class is oversampled by joining the minority class’s 5 nearest neighbors in high dimensional space. This means SMOTE operates on the feature space, rather than the data space. We generated these synthetic samples by taking the difference between the feature vector (sample) and one of its 5 nearest neighbors, multiplying by a random number between 0 and 1, and adding it to the sample under consideration. This creates a new “synthetic” sample along a random point on the line in high dimensional space. Doing this forces the generalization of the minority class’s decision region and causes the classifier to create larger, less specific decision regions [21]. For our analysis, we used SMOTE from Python’s imbalanced learn package. The resulting dataset becomes balanced with 260 individuals for each diagnosis.

## 3. METHODS

We first developed a RF classifier using blood microarray gene expression data from 744 participants in the ADNI cohort. Then, UMAP was used for dimensionality reduction to validate the DEGs, and HDBSCAN was used to identify subclusters of AD and reveal important DEGs in each subcluster. After that, GFCI was used to find potential causes of AD within the DEGs. Finally, GSEA was used to identify relevant biological processes and pathways enriched in each phenotype (Figure 1A).

**Figure 1.**
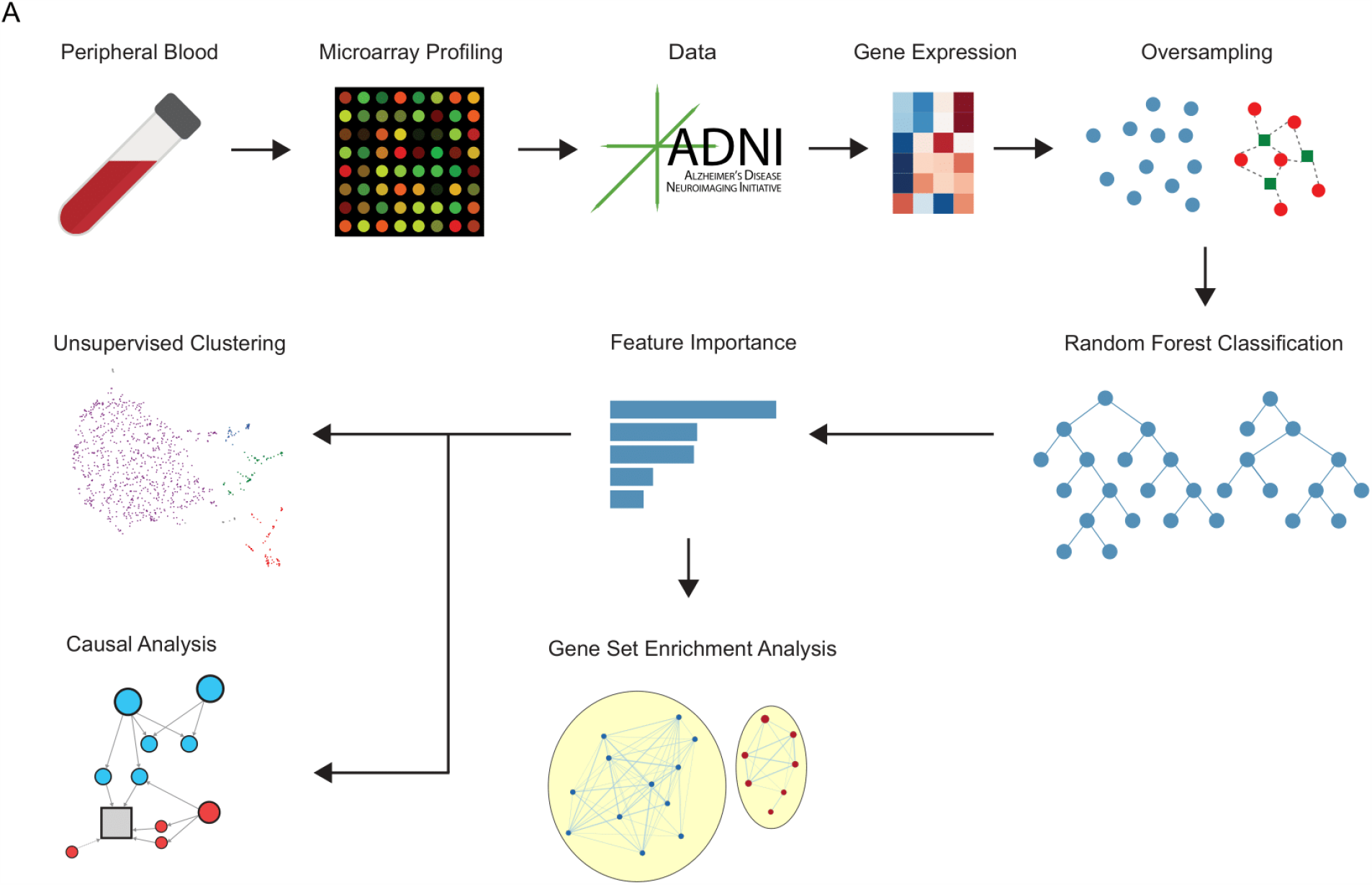
Workflow. (A) Workflow shows data acquisition, preprocessing, and analysis.

### 3.1 Random Forest Classification

RF is an ensemble learning algorithm that constructs a forest of binary decision trees learned by randomly sampling from the training set. These trees (the forest) can then be applied to a given sample to generate a class probability that reflects its similarity to a given class of the training set. We chose to use a RF classifier as it generally avoids overfitting because of the low correlation between trees. In this case, we used supervised learning to train the RF classifier on patient’s blood gene expression values and predict a clinical diagnosis of CN, EMCI, LMCI, or AD. The model was evaluated on the four categorical classes and two binary classes where the two classes represented AD compared to control (CN, EMCI, LMCI). For our purposes, we used an implementation of RF from Python package skLEARN with 1000 estimators (trees).

We used RF classification to (1) predict if an individual had AD using their gene expression profile and (2) to obtain a ranked gene list of the most important features in classifying AD. First, we performed a feature selection step for the RF classifier. This step selected the most informative features from all 48,157 probe transcripts. To do this, we first trained an ‘‘outer’’ RF classifier on the dataset. We trained a forest with 1000 trees, using 80% of the individuals as a train set and the remaining 20% as a validation set. Based on the feature importance scores of the ‘‘outer’’ RF classifier, we then selected the top 250 most informative genes for training of the ‘‘inner’’ classifier. This ‘‘inner’’ classifier was further evaluated through 5-fold cross-validation. During cross-validation, the dataset was split into five smaller sets of which four were used to train a RF classifier that was tested on the remaining set. This “inner” classifier was then used to build a final RF classifier to compare to other popular machine learning models using only the top 25 most informative genes. We compared RF to neural network, support vector machine, naive bayes, and logistic regression. These models were implemented from Python package skLEARN with default parameters and compared via 5-fold cross-validation.

Finally, we obtained a ranked gene list of the most important features in classifying AD using feature importance scores from the RF classifier. After training and validation, we estimated which variables were important in the classification using the Gini impurity metric. RF’s locally optimal condition is chosen based on Gini impurity. When training a tree, we computed how much each feature decreased the weighted impurity. Then, we averaged the Gini impurity decrease from each feature in the forest and assigned each feature a feature importance score based on this decrease. The sum of all the feature importance scores is 1 and genes with higher feature importance scores play a greater role in the decision trees of the RF model. To validate the feature importance independently we also calculated the Wilcox score of each gene and compared the gene’s p-value with its rank. In general, the higher a gene was ranked in the RF classifier the lower its p-value.

### 3.2 Unsupervised Clustering

We employed uniform manifold approximation graphs (UMAP) for dimension reduction and unsupervised clustering. UMAP is a manifold learning technique for dimension reduction and is competitive with tSNE for visualization quality while preserving global structure [22]. We used UMAP to visualize the similarities among individuals within each class and identify sub clusters of AD. We selected the set of top 100 DEGs from the 250 gene RF classifier. This is because the first 100 DEGs contribute most to the accuracy of the RF model meaning the rest likely provide the little additional information. For initial dimensionality reduction, we used Principal Component Analysis (PCA) and selected the top 20 PCs based on the elbow plot. For UMAP visualization we used the package UMAP-learn in Python with 30 nearest neighbors, a minimum distance of 0.01, and 2000 epochs [75]. We also visualized the connectivity in the manifold by simplifying the intermediate topological representation of the approximate manifold down to a weighted graph. This helped us further visualize the relationships between clusters and individuals.

To diagnose the validity of UMAP we utilized three diagnostic graphs: PCA, local dimension, and local neighborhoods. The PCA diagnostic takes the first 3 PCs, which preserve global structure, and convert the coordinates of each point into an RGB description of a color. By projecting the colors onto the UMAP embedding we can see if UMAP successfully captures the global structure. For local dimension, each data point’s local dimension should ideally match the embedding dimension. In practice, a high local dimension represents points that UMAP has difficulty embedding. Thus, the embedding tends to be more accurate in regions where the points have consistently lower local dimension. Finally, we visualized preserved local neighborhoods in terms of the Jaccard index. A higher Jaccard Index means the local neighborhood has been more accurately preserved. All three diagnostics support accurate global structure and embedding of AD individuals.

Finally, we used HDBSCAN for unsupervised clustering to identify subclusters of AD and reveal important DEGs in each subcluster. We chose HDBSCAN as it is a hierarchical clustering algorithm with the ability to cluster data of varying shapes and densities. We used the Python implementation of HDBSCAN on UMAP space with a minimum cluster size of 30 and 5 minimum samples. Heatmaps with dendrograms were drawn to identify differences between clusters. The gene expression of top marker genes from each cluster was projected onto UMAP to identify cluster specific genes.

### 3.3 Causal Analysis

We used the Greedy Fast Causal Inference (GFCI) [77] algorithm to perform our causal analysis. For the causal search, we used continuous gene expression values for the 100 DEGs selected by the RF and discrete diagnosis values as our data. The diagnosis values were either binarized (0: AD and 1: Non-AD) or categorical (0: AD, 1: LMCI, 2: EMCI, and 3:CN)., We used the Tetrad interface [23] to implement GFCI with an alpha value of 0.05 to analyze potential causes within and across the DEGs and the phenotype.

The GFCI algorithm requires [24]: (1) The causal Markov condition holds true. This happens if a variable is independent of all of its causal non-descendants when conditioned on its causal parents [25]. (2) The causal faithfulness condition holds true. This condition states that all causal hypotheses are probabilistically dependent [26]. (3) There is no missing data. (4) There is no selection bias. (5) The measured variables consist of no feedback cycles. We assume the causal Markov and causal faithfulness conditions hold true for our data, and the rest of the conditions are met.

Our implementation of the GFCI algorithm runs based on a Bayesian search algorithm that searches the space of directed acyclic graphs (DAGs). In particular, it starts with an empty DAG and then performs a search where nodes are connected until the Bayesian score cannot increase with the addition of an edge. Then, a backward-propagation stepping search is performed where edges are removed until the score can no longer increase [27]. Finally, a series of conditional independence tests remove edges between two variables that are evaluated to be independent [28]. We chose to implement the GFCI algorithm because, unlike numerous other search algorithms, it does not operate under the assumption that there are no latent confounders and outputs when there is a possibility of a latent confounder [30].

### 3.4 Enrichment Analysis

Gene Set Enrichment Analysis (GSEA) [54] was used for functional analysis of the top 2000 DEGs identified by the RF classifier. We chose the top 2000 features because they have relatively high feature importance scores from the RF classifier (Figure 3C).

We used the oversampled dataset with 260 samples for each diagnosis (AD, LMCI, EMCI, CN) and polarized the dataset by only selecting the samples that were part of the AD and the CN groups. This was done in hopes of avoiding confounding and allowing easier enrichment in one phenotype over the other.

We ran our analysis on the C5 Gene Ontology (GO) gene sets found in the Molecular Signature Database (MSigDB). We used Benjamini-Hochberg adjusted p values (q values / FDR) for multiple testing correction [55]. Gene sets with a q<0.25 and nominal p<0.01 were considered significantly enriched. To screen for the upregulated and downregulated gene sets, we ran enrichment analysis with the following parameters: 1000 permutations, weighted enrichment statistic, Signal2Noise ranking metric, a maximum size of 500, and a minimum size of 3.

GSEA also provides a ranked genes list which ranks the genes according to this Signal2Noise metric, which measures a gene’s correlation with the phenotype. A positive value indicates correlation with AD and a negative value indicates correlation with CN.

Further, we conducted analysis to evaluate how these significantly enriched gene sets interacted with each other. We first created an enrichment map. The jaccard coefficient was used to measure the similarity between the gene sets. Gene sets with a jaccard coefficient<0.25 were considered correlated and connected via an edge in the Enrichment Map. We visualized the Enrichment Map using Cytoscape [56]. Then, AutoAnnotate [57] was used to cluster the gene sets and annotate each cluster with a label. The MCL cluster algorithm was used to cluster the gene sets and the WordCloud adjacent words algorithm was used to label each cluster. We then manually adjusted each annotation to ensure it represented the biological theme and removed clusters with less than two gene sets.

Finally, leading edge analysis was conducted using the GSEA software on all gene sets considered significantly enriched. Leading edge analysis works by analyzing which genes contributed most to the enrichment score (ES) of a given gene set’s leading edge [58]. This was done in order to determine which genes have the biggest impact on the enrichment of multiple gene sets.

## 4. RESULTS

For the 744 ADNI samples, the average chronological age was 73.14, and the ratio of females was 45.16%. Both ages and gender distributions did not differ significantly between AD and CN samples (Supplementary Table 3A). Additionally, there was no statistically significant difference between gender and ages between AD and non-AD (Control) samples (Supplementary Table 3B).

### 4.1 High Sensitivity Random Forest Model is able to distinguish AD from Controls

We created a RF classifier to classify AD from non-AD individuals and to identify the most important genes related to AD. We first utilized an initial feature selection step consisting of an “outer” RF classifier that selects the top 250 features based on feature importance. We chose the top 250 features because they capture a majority of the feature importance. Next, we created an “inner” RF classifier using the top 250 genes selected by the “outer” RF. After initial feature selection, 5 fold cross-validation of the 250 gene classifier achieved an accuracy of 99.04%. The RF classifier also achieved 99.2% sensitivity detecting AD and 98.97% specificity detecting non-AD individuals (Figure 2B and Table 1B). The high sensitivity of the RF classifier supports the possibility of an accurate and minimally invasive screening tool for AD.

**Figure 2.**
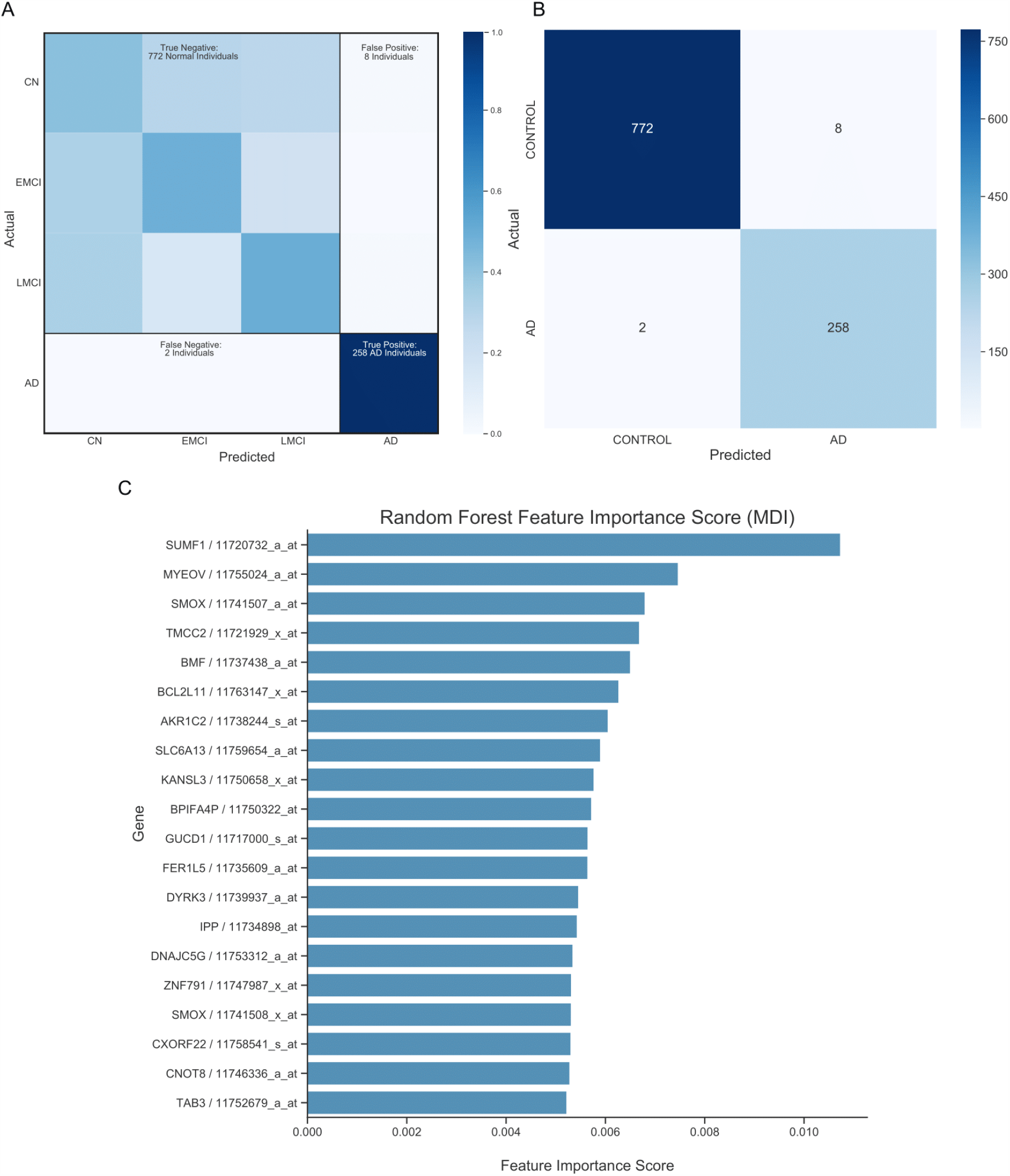
Random Forest Classification. (A)Heatmap depicts results of a 5-fold cross-validation of the “inner” RF classifier comprising 4 classes for the 250-gene classifier. Individuals that fall on the diagonal are classified correctly (59.7% of individuals). Individuals that do not fall on the diagonal are mis-classified as a different diagnosis. (B)Confusion Matrix depicts results of a 5-fold cross-validation of the “inner” RF classifier comparing AD to Control (CN, EMCI, LMCI) for the 250-gene classifier. The sensitivity of detecting AD (true positive rate) is 99.2%. The specificity of detecting AD (true negative rate) is 98.97%. This data supports the high accuracy of the machine learning classifier. (C)Barplot shows the top 20 genes of the “inner” RF classifier ranked based on feature importance score.

**Figure 3.**
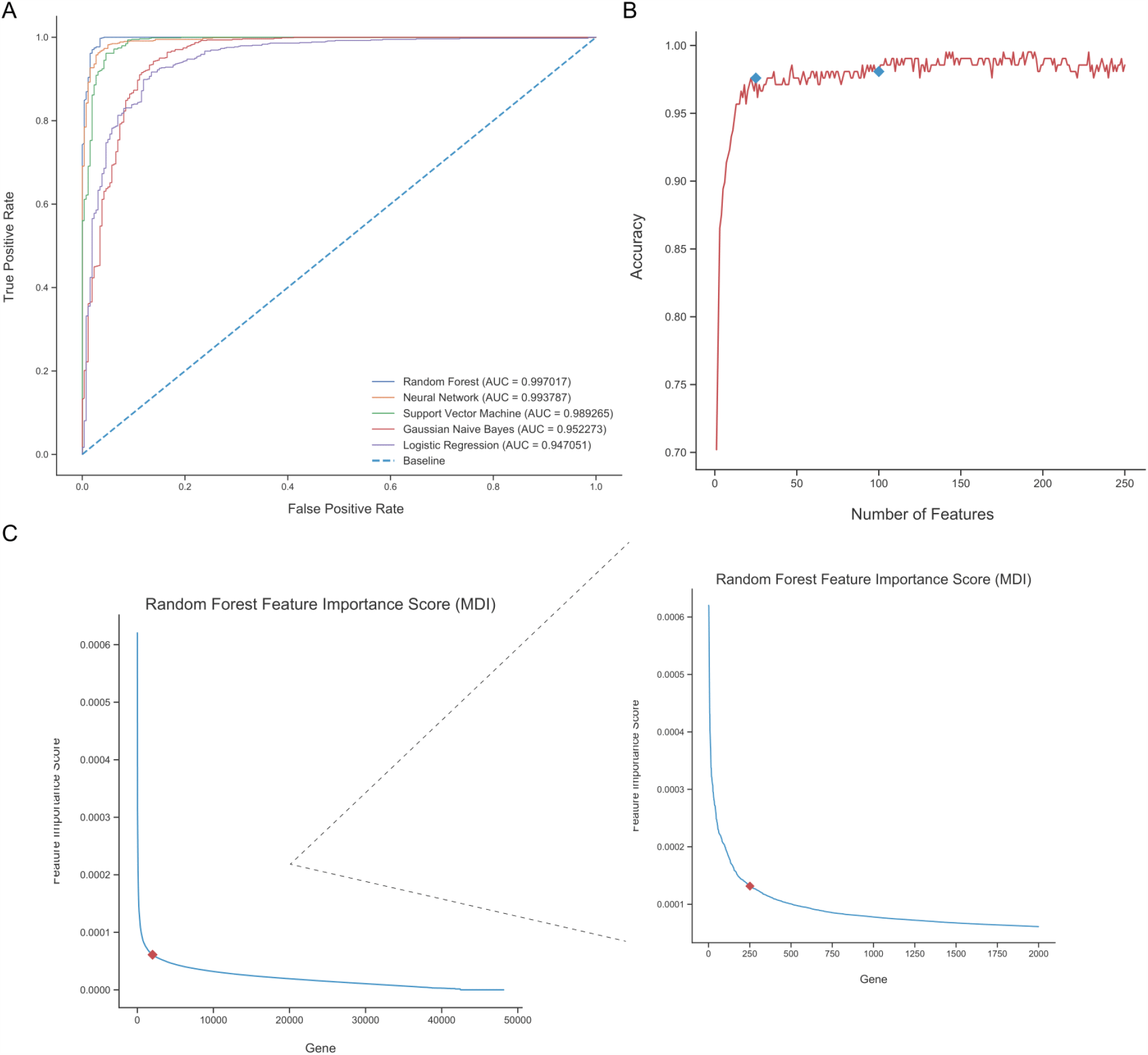
Random Forest Metrics. (A)Receiver operating characteristic curve for RF, Neural Network, Support Vector Machine, Gaussian Naive Bayes, and Logistic Regression on the top 25 gene’s in feature importance. (B)Line plot shows impact of feature number on RF accuracy. At 100 features the RF model reaches a high accuracy while avoiding overfitting. Prior to 25 features the RF model underfits and achieves drastically lower accuracy. (C) Line plot (Left) shows each gene’s feature importance score in the “outer” RF classifier. The top 2000 features capture a majority of the importance in the model. Among the 2000 features (Right) the top 250 features capture a majority of the feature importance.

A heatmap depicts the results of 5-fold cross-validation of the “inner” RF classifier composed of 4 classes (Figure 2A). The first three classes are CN, EMCI, or LMCI. The gene expression signatures from peripheral blood are not only able to strongly differentiate AD from non-AD individuals but are also able to differentiate CN, EMCI, and LMCI, albeit at a much lower accuracy. The RF classifier achieves an accuracy of 59.7% when differentiating between all 4 diagnosis groups (Figure 2A). This suggests changes in gene expression also occur during MCI that differ from both AD and CN.

We ranked the top 100 features from the RF classifier based on feature importance. Figure C shows the top 20 genes contributing to the RF classification of the individuals. Sulfatase Modifying Factor 1 (SUMF1) appears as the most important feature in the RF classifier. This means SUMF1 plays the largest role in the decision trees of the RF model and may be a significant risk factor of AD. Additionally, SMOX appears as the third most important feature in the RF classifier. This is key as SMOX is a gene that consistently surfaces in our analysis.

Finally, we compared the performance of our RF model to other popular machine learning models, namely: Neural Network (NN), Logistic Regression (LR), Gaussian Naive Bayes (GNB), and Support Vector Machine (SVM) models. The 250 gene RF classifier achieved an AUC score of 0.99. However, this score is quite similar to other models trained on the same 250 genes with NN and LR actually achieving a higher AUC (Supplementary Figure 1C). RF is robust to overfitting compared to other models so we hypothesized that the high AUC of other models may be from overfitting the data. This becomes clear when comparing 25 gene models. We chose to use 25 genes as the RF classifier is able to maintain relatively high accuracy at 25 genes (Figure 3B). Less features, however, substantially impacts the accuracy of the other models. When comparing the 25 gene models, RF outperforms all other models with an AUC of 0.997 (Figure 3A). The ability of the RF classifier to maintain a high AUC with a low number of features suggest the possibility of an accurate and minimally invasive screening tool for AD with a small, robust set of genes.

### 4.2 Unsupervised Clustering reveals 3 distinct AD populations

After RF classification, we performed unsupervised clustering to identify subclusters of AD and reveal important DEGs in each subcluster. For initial dimension reduction we performed Principal Component Analysis (PCA) and selected the top 20 principal components for further dimension reduction. Even in PCA space AD individuals are relatively separated from EMCI, LMCI, and CN (Figure 4C). This means at a global scale AD individuals have distinct gene expression profiles. After initial PCA dimension reduction UMAP was employed to further reduce dimensionality. UMAP is able to clearly differentiate AD from non-AD individuals (Figure 4D). Dimensionality reduction techniques show blood microarray gene expression of AD individuals can be separated from normal individuals. The individuals closer to AD clusters have genetic signatures similar to AD individuals and may be at risk for AD. In fact, the RF classifier was more confident (higher classification probability) when classifying AD individuals that separated from the main population. This validates the genes identified by the RF classifier as viable biomarkers for AD screening and detection.

**Figure 4.**
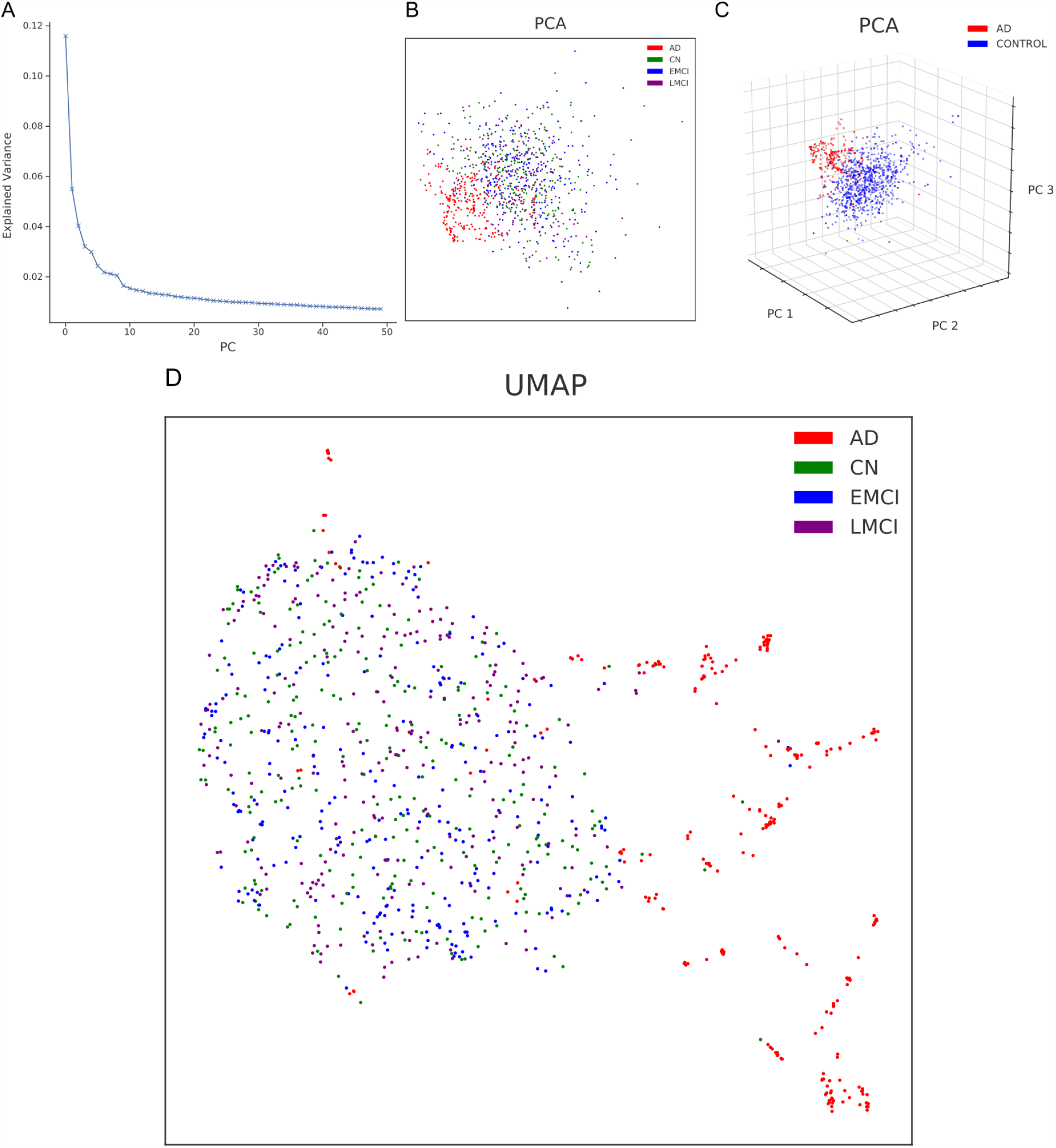
Dimension Reduction. (A)Elbow plot shows the explained variance of each principal component. We chose the top 20 PCs for downstream analysis. (B)PCA visualization of 744 individuals (points), with similar individuals positioned closer together. Points are color-coded by diagnosis. (C)Three dimensional PCA visualization of 744 individuals (points). Points are color-coded by binarized diagnosis either AD or Control (CN, EMCI, LMCI). (D)UMAP visualization of 744 individuals (points), with similar individuals positioned closer together. Points are color-coded by diagnosis.

We then used HDBSCAN to cluster individuals on UMAP space. Unsupervised clustering results in three distinct clusters of AD-like individuals (Figure 5A and 5C). The connectivity of the manifold suggests that the AD clusters are still connected to the Control cluster but are substantially more interconnected (Supplementary Figure 2A). Specifically, AD3 is less connected with the main population while AD1 and AD2 are relatively connected with each other. To explore the relationships between these clusters we identified DEGs among the clusters. Gene expressions were ranked between the groups based on variance and a cluster heatmap drawn for the 20 genes with the highest variance. The heatmap shows distinct gene expression profiles that distinguish AD clusters from Control and each other. Spermine Oxidase (SMOX) appears in unsupervised clustering and as the third most important feature in the RF classifier (Figure 2C). This means SMOX likely plays a strong role in AD as it appears as an important feature in both unsupervised and supervised methods.

**Figure 5.**
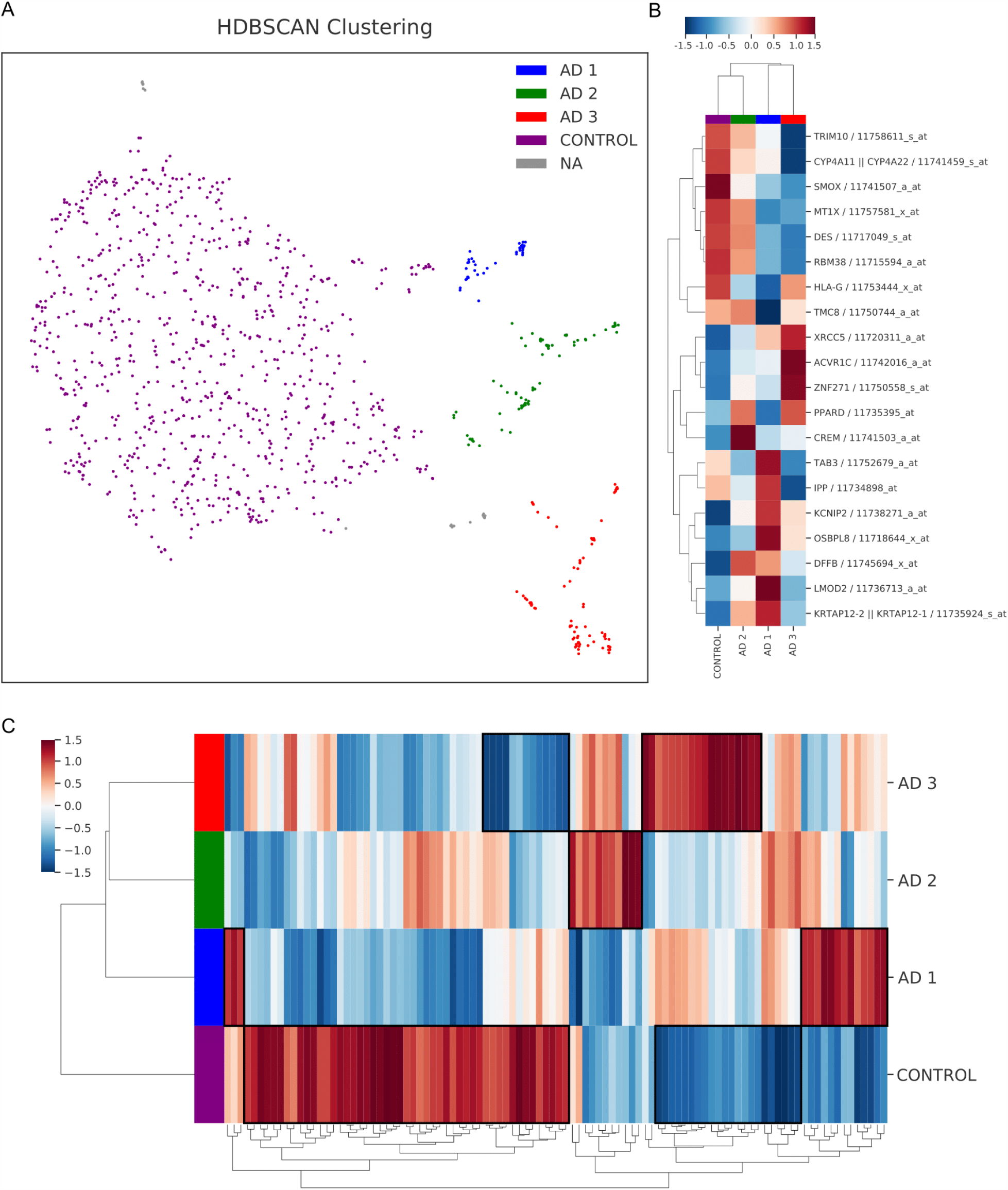
Unsupervised Clustering. (A) UMAP visualization (as in Figure 4D) overlaid with clusters determined by unsupervised clustering. HDBSCAN clustered the individuals into 3 AD clusters and a control cluster. (B)Heatmap shows the expression of the top 20 differentially expressed genes (rows) determined by Wilcoxon rank sums test among the HDBSCAN-defined clusters (columns). The rows and columns are grouped by hierarchical clustering. (C)Heatmap shows the expression of all 100 genes used in unsupervised clustering with differentially expressed genes as columns and HDBSCAN-defined clusters as rows. The rows and columns are grouped by hierarchical clustering. The figure confirms the heterogeneity of AD and the three AD subclusters identified.

### 4.3 Causal Analysis identifies potential causal factors and key hub genes

Finally, we conducted Causal Analysis on the DEGs selected by the RF using GFCI. This was done to find potential causes of AD. The resulting DAG is visualized in Cytoscape through an edge-weighted spring embedded layout. In the causal graph two major hub genes emerge. First, the Major Histocompatibility Complex, Class II, DP Beta 1 (HLA-DPB1) gene emerges as the largest node. It is a central hub gene because out of the 100 DEGs, HLA-DPB1 has potential direct causal relationships with 75 of them. Additionally, HLA-DPB1 connects to the phenotype (AD diagnosis) through SMOX. HLA-DPB1 also indirectly influences the second-largest node, CYPA411 which is then connected to AD through SUMF1 and SMOX (Figure 6A).

**Figure 6.**
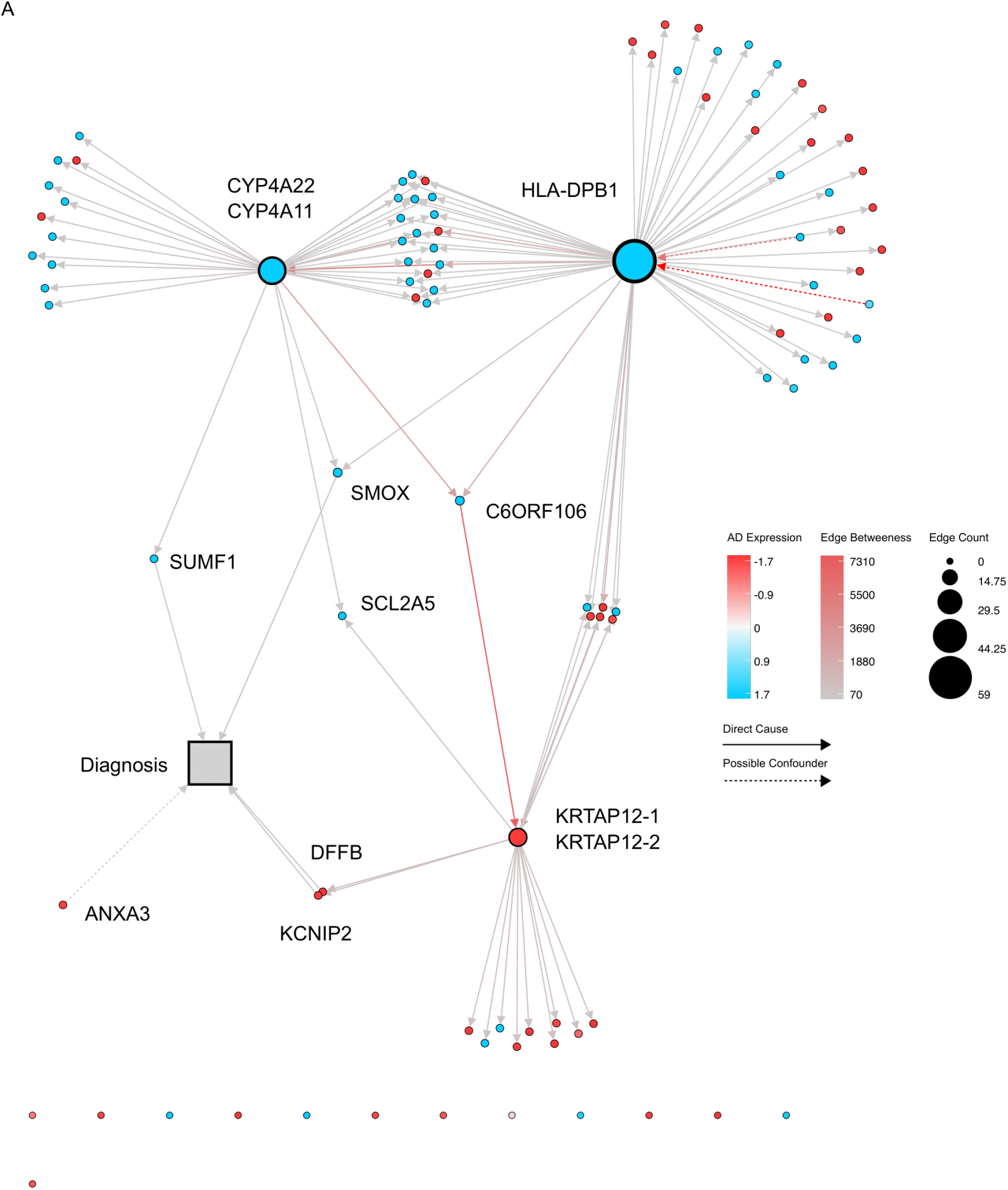
Causal Analysis. (A) Directed Acyclic Graph from GFCI causal analysis is visualized in Cytoscape through an edge-weighted spring embedded layout. Edge betweenness centrality is an edge centrality index. A high betweenness score means the edge is crucial to preserving node connections.

Due to the high connectivity of HLA-DPB1, it is likely this gene plays a significant role in the molecular mechanisms underlying AD through downregulation. Second, CYP4A11 emerges as the next largest node having an edge count of 42. Although CYP4A11 does not have a direct causal relationship with the phenotype, it is a hub gene and may play some underlying role.

In addition to the two hub genes, four direct connections with the phenotype (AD diagnosis) also emerge in the causal graph. SUMF1, SMOX, KCNIP, and DFFB all have causal relationships with AD with no latent confounder (Figure 6A). SUMF1 and SMOX have been identified as AD-related genes in every other method we have used. This essentially validates the causal graph and shows those with a direct connection to the phenotype likely are correlated with AD.

### 4.4 Gene Set Enrichment Analysis reveals enriched biological processes

First, the genes ranked by the Signal2Noise metric corroborate key AD genes identified by previous methods. For example, SUMF1, SMOX, and DFFB all emerge as having some of the highest absolute values of this metric (Figure 7A) (Table 1).

**Figure 7.**
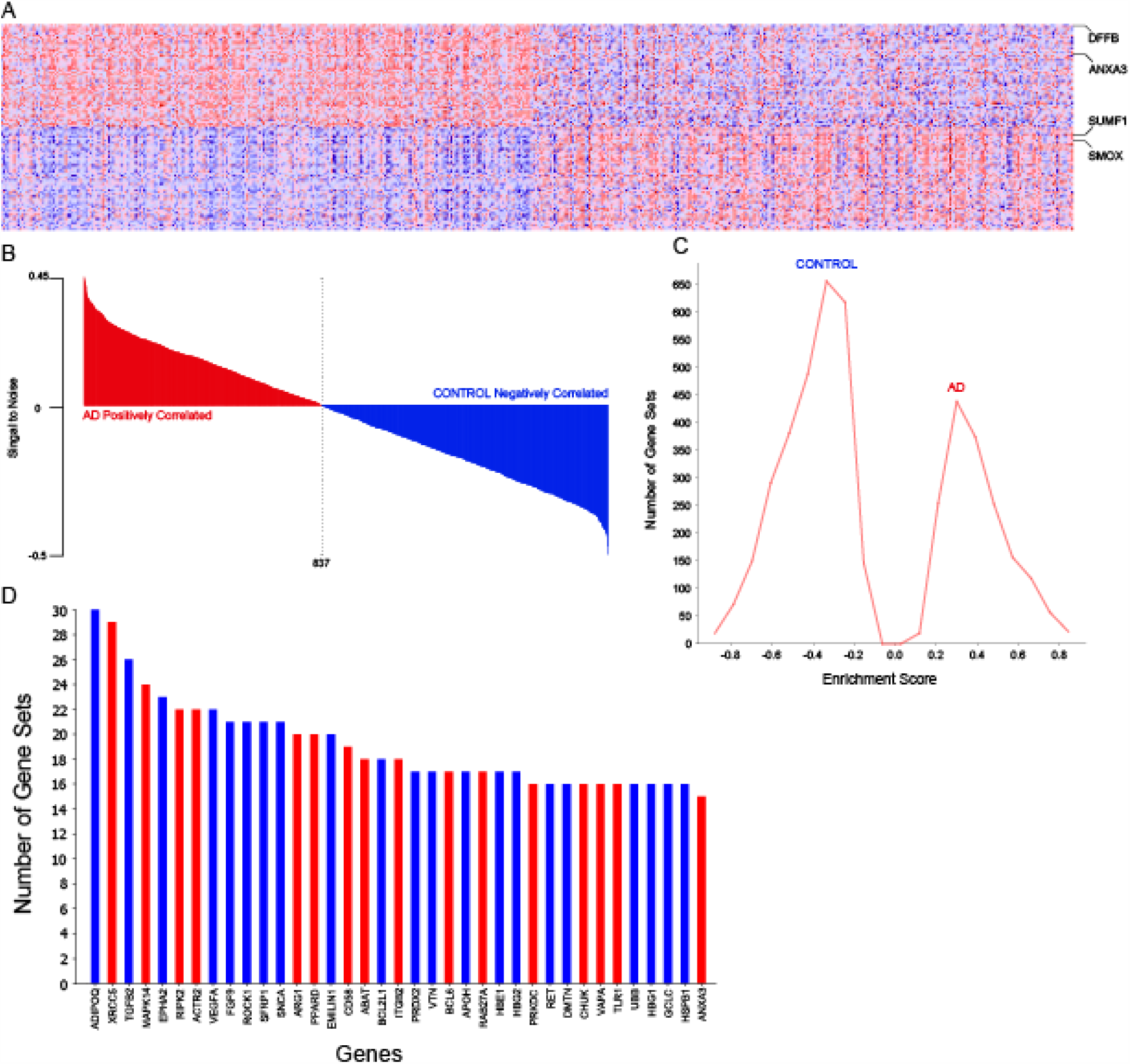
Gene Set Enrichment Analysis. (A)Heatmap depicts how the top 50 genes upregulated in AD and the top 50 genes downregulated in AD (according to the ranked genes list) differ among the two classes (AD and CN) (B)Ranked Gene List Correlation Profile shows how the metric for ranking genes (Signal2Noise) changes based on the position a gene is in the ranked genes list. This supports that genes towards both extremes of the ranked genes list are highly differentially expressed. (C) Global Enrichment Score Histogram depicts number of gene sets across enrichment scores.

Further, we ran GSEA on the top 2000 DEGs to find which GO term gene sets were suggested to play a role in AD based on our DEGs. Among the enriched gene sets, biological processes related to AD pathogenesis emerge, including processes related to Cell Activation in the Immune System, Glycosaminoglycan (GAG) Binding, vascular dysfunction, oxidative stress, and the neuron apoptotic process emerge (Figure 8A). More details regarding individual gene sets can be found in Supplementary Figure 3.

**Figure 8.**
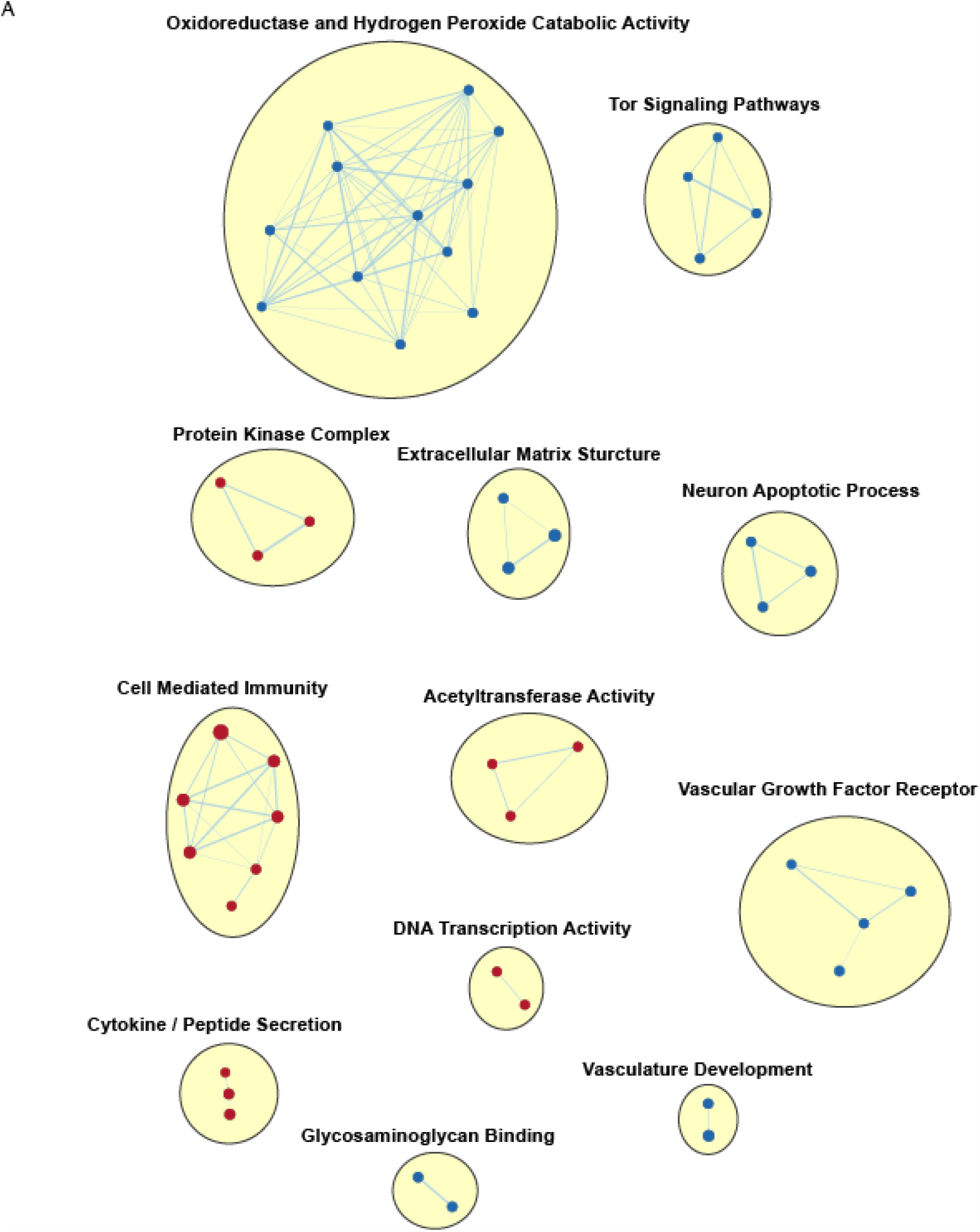
Enrichment Map. (A)Enrichment Map representation of the GSEA results obtained for AD versus CN. Red represents enrichment in AD, whereas blue represents enrichment in CN; color intensity is proportional to enrichment significance. Node size represents gene set size and edge width represents number of shared genes. Clusters of functionally related gene-sets were labeled using AutoAnnotate and manually adjusted to match the overall biological theme.

Cell Activation in the Immune System is suggested to play a role in AD pathogenesis. Researchers have shown that Aβ-reactive CD4 T cells are able to effectively target Aβ plaques in the brains of APP-transgenic mice [60]. Although other studies have not connected HLA-DPB1’s function to specifically AD, HLA-DPB1 is key in the recognition and activation of CD4 T-cells[59]. Thus, we hypothesize downregulation of HLA-DPB1 (a hub gene in our causal analysis) contributes to AD pathogenesis through lower activation of Aβ-reactive CD4 T cells which may contribute to the removal of these toxic plaques.

GAG binding is another key pathway through which our DEGs play a role in AD. Glycosaminoglycans have already been proven to play a role in AD as the sulfate moieties of GAGs play a critical role in the binding to Amyloid betas and enhance Amyloid beta fibril formation [61]. This also reflects the role of SUMF1, the top gene in the RF classifier. The SUMF1 gene activates sulfatases that play a role in many mechanisms of lysosomes including degradation of GAGs and macromolecules [31]. SUMF1 is downregulated in AD, so we hypothesize it progresses AD by decreasing the degradation of the GAGs.

Another biological process that emerged in GSEA is vascular dysfunction. Some of the key pathways related to vascular dysfunction, including vascular growth factor receptor and vasculature development, are downregulated in AD. These pathways have been previously linked to AD pathogenesis. In fact, vascular dysfunction has been potentially shown to precede hallmarks of AD by contributing to parenchymal amyloid deposition, neurotoxicity, and glial activation [62]. This is tightly linked with CYP4A11, a major hub gene in our causal analysis. CYP4A11 produces 20-HETE and is directly involved in vascular disfunction [35]. CYP4A11 is downregulated in AD, so we hypothesize it progresses AD through impaired cerebral autoregulation.

Finally, the neuron apoptotic process emerges as a key pathway in GSEA. Plaques and neurofibrillary tangles are strongly associated with massive neuronal apoptosis, especially in the cerebral cortex and hippocampus. This neuronal apoptosis has been evident in AD onset and progression [75]. Numerous genes from the causal analysis and leading-edge analysis relate to apoptosis, the top being DFFB.

## 5. DISCUSSION

Functional genetic biomarkers linked to AD can greatly increase timely diagnosis and intervention, especially in the predementia phase. Cerebrospinal fluid (CSF) biomarkers correlate well with AD but lumbar punctures are invasive. This means CSF is not suitable for use in large scale screening of AD. Similarly, positron emission tomography (PET) imaging of amyloid beta protein in the brain often reveals accurate prognosis, but PET imaging is expensive and impractical. On the other hand, blood biomarkers linked to AD are easily accessible and inexpensive allowing for large scale early screening.

This study identified and evaluated a blood gene expression diagnostic classifier of AD that could distinguish people with AD from normal control subjects. Cross-validation results show the “inner” RF classifier achieved an accuracy of 99.04% and a sensitivity of 99.2% (Table 2B). The high accuracy and sensitivity of the model supports the possibility of powerful and minimally invasive tools for early clinical screening and AD prevention. This study also identifies two genes that have consistently been chosen as relating to AD across all our methods: SUMF1 and SMOX.

To our knowledge, limited research has directly identified SUMF1 as a risk factor of AD. However, literature suggests that SUMF1 plays a key role in other neurodegenerative disorders. Expression of SUMF1 in brain and visceral tissues results in activation of sulfatases. In severe multiple sulfatase deficiency (MSD), mutations in the SUMF1 gene of astrocytes result in lower global sulfatase activity and defective degradation of autophagosomes [45]. Specifically, SUMF1−/− astrocytes have defective autophagy pathways [73]. This is prominent in both MSD and neurodegenerative diseases as impaired autophagy increases cytoplasmic protein aggregation [45]. In fact, dysfunction of the autophagy pathway in astrocytes has been linked to the accumulation of misfolded proteins in AD [74]. Our results theorize a prominent role of sulfatases through astrocyte dysfunction and impaired autophagy in AD neurodegeneration.

Also, few existing literature identifies a correlation between SMOX and AD. Even among the literature that identifies a correlation, they do not expand upon SMOX’s role as a potential AD-related gene [79][80]. We hypothesize downregulation of SMOX contributes to AD thorough oxidative stress. Oxidative stress is a key hallmark of AD [64]. GSEA results suggest that this occurs through a downregulation in the oxidoreductase/hydrogen peroxide catabolic process which is the largest cluster in our enrichment map. SMOX catalyzes the oxidation of spermine [33] which results in the generation of hydrogen peroxide (H2O2) [65]. The imbalance of oxidants and antioxidants, especially reactive oxygen species (ROS) such as H2O2, is a key cause of oxidative stress. Severe downregulation of SMOX could cause this imbalance and progress AD pathogenesis through oxidative stress. Already, the brains of patients from AD have been identified as suffering from a significant extent of oxidative damage where brain proteins are oxidized by free radicals, affecting enzymes critical to neuron and glial functions [66][67]. In fact, abnormal oxidation of the tau protein has been reported as a key post translational modification tau undergoes to aggregate into paired helical filaments (PHFs), which make up neurofibrillary tangles [68][69][70]. We hypothesize downregulation of SMOX may contribute to oxidative stress and the resulting accumulation of AD pathological hallmarks.

Additionally, our implementation of the GFCI algorithm further reveals potential AD-related genes. These genes include the hub genes in our causal graph (HLA-DPB1 and CYP4A11) and genes that had a direct causal connection with AD with no latent confounder (SUMF1, SMOX, and AD). Although true causality cannot be established through our causal analysis, it does provide potential and hypothetical causes of AD for further research and experimentation. Additionally, the genes above could all be corroborated in multiple methodologies as being related to AD. Rather than definitively declare a genetic cause, we hope to provide genes of interest for future research that can be done to validate findings in the lab.

The results of the causal analysis are already partially validated by literature searches, such as identifying DFFB and HLA-DPB1 as AD-related genes. DFFB has a direct causal connection with AD and has the top rank in GSEA’s ranked genes list. DFFB codes caspase-dependent DNase which triggers apoptosis through DNA fragmentation and chromatin condensation [71]. Previously, experimental AD models in rats have shown that glutamate antibodies, which repress DFFB, stabilize AD processes and decrease the intensity of the apoptotic death of neurons and glial cells [72]. This is consistent with our results which suggest, inversely, that upregulation of DFFB contributes to AD pathogenesis. We validate these findings in addition to suggesting that DFFB may play an underlying role in AD in homo sapiens, not just rats.

Additionally, previous studies have identified copy number variants (CNVs) in HLA-DPB1 in both the blood and brain tissue correlating with AD. HLA-DPB1’s previous association with AD helps to validate our model. Although HLA-DPB1 has been identified as a CNV in previous studies, it has not been suggested to change in expression. Our study suggests HLA-DPB1 is downregulated and may play a role interacting with other genes. We hypothesize HLA-DPB1 plays a role in AD through its activation of CD4 T-Cells [78].

In addition to confirming previously identified genes, our causal model provides new genes of interest. CYP4A11 is a major hub gene in our causal analysis and a member of the CYP family, which produces 20-HETE [35]. 20-HETE plays a critical role in the vasculature through vasoconstriction, cerebral autoregulation, inflammation, and maintaining blood-brain barrier integrity [63]. Specifically in AD, there is a reduction in the number of micro vessels, vascular smooth muscle cells (VSMCs) and flattening of endothelial cells [46], suggesting AD may be linked to impaired cerebral autoregulation. The vessel’s ability to autoregulate with the rise and drop in blood pressure is achieved mainly through the myogenic response which is enhanced through metabolic activators such as 20-HETE. Thus, downregulation of CYP4A11 in AD leads to reduced 20-HETE production and autoregulation impairment, which is a symptom of AD. Our study advances this with the suggestion of CYP4A11 as a potential risk factor for AD through its role in the creation of 20-HETE [35]. This is appropriate given our gene expression samples are collected from the peripheral blood.

Next, unsupervised clustering using UMAP and HDBSCAN yielded interesting results. Clustering of AD individuals did not result in one cluster. Instead, the results of clustering suggest three separate groups of AD individuals with unique gene expression signatures. This suggests AD may manifest in multiple different genetic pathways. This also suggests different causes of AD, further supporting AD’s heterogeneity.

The present study has some limitations regarding data and methodologies. First and foremost, our gene expression samples come from PBMCs and not tissue samples in the brain. This limits our claim that key genes identified in this study may play an underlying role in AD which is a neurological disorder. However, this does not detract from the main purpose of this paper which is to aid in the screening of AD. We provide a model and key genes in hopes that they could be used to develop a robust genetic screening process to detect AD early. Second, selecting genes based on the RF feature importance may remove correlated features. When the data has two features that are correlated, the RF model can not differentiate between them. Any of these correlated features during training and subsequent classification can be used with no concrete preference of a certain feature. However, this effect is reduced due to random sampling at each node creation. Next, the ADNI data is not equally represented and thus imbalanced. To overcome the imbalanced data problem, we used an oversampling technique called SMOTE. Although this balances the dataset, it calls into question the efficacy of the algorithm and its possibility of producing an overfitted model. Additionally, causal analysis can not establish true casualty as it only provides potential and hypothetical causes of AD for further research and experimentation. However, genes identified provide opportunities for further analysis in their potential role of AD. Finally, filtering pathways based on a nominal p-value<0.01 and an FDR<0.25 in our GSEA analysis does mean it is more likely for there to be false-positives due to multiple-hypothesis testing; however, as stated earlier, our DEGs’ function and how they play a role in AD is not the key takeaway.

## 6. CONCLUSION

The detection of AD at the earliest possible stage is key to support advances in AD intervention, prevention, and treatment. Our analysis provides a high-accuracy RF classifier that uses gene expression from peripheral blood to differentiate AD from normal controls with SUMF1 emerging as the most important feature, suggesting a role of sulfatases and astrocyte dysfunction in neurodegeneration. Unsupervised clustering analyzed relationships between DEGs and identified downregulated SMOX expression potentially playing a role in AD through a breakdown in regulation of polyamines. Finally, causal analysis using GFCI to analyze the DEGs’ potential causal relationships with AD identified HLA-DPB1 and CYP4A11 as hub genes and DFFB as a potential cause of AD. HLA-DPB1 has been identified previously as a CNV associated with AD but our study suggests downregulation on the expression level. Additionally, repression of DFFB has been shown to stabilize the effects of AD, including neuron death, in rat models, but our results suggest it may play a role in homo sapiens as well. The genes identified in this study may play an underlying role in AD and neurodegeneration through their respective functions.

This study further advances genetic screening of AD through a robust RF classifier of peripheral blood gene expression and provides functional genetic biomarkers and potential gene targets for future experimentation. In addition, the three clusters of AD individuals identified suggest AD may manifest in multiple different genetic pathways that can be further explored. Finally, integrating blood measures like proteins and metabolites along with gene expression may further improve biomarker identification and classification accuracy.

## 7. ACKNOWLEDGMENTS

We sincerely thank Dr. David Boone for his insights during the conception of this study and his ongoing technical support. We thank ADNI for providing data for this study.

## 9. DATA AND CODE AVAILABILITY

Microarray gene expression data can be accessed from the ADNI database (adni.loni.usc.edu). Python notebooks written for data analysis, random forest, unsupervised clustering, and other analyses are shared via GitHub (https://github.com/andrewni4313/geneticAD2020).

## 11. SUPPLEMENTARY TABLES

**Supplementary Table 1.**
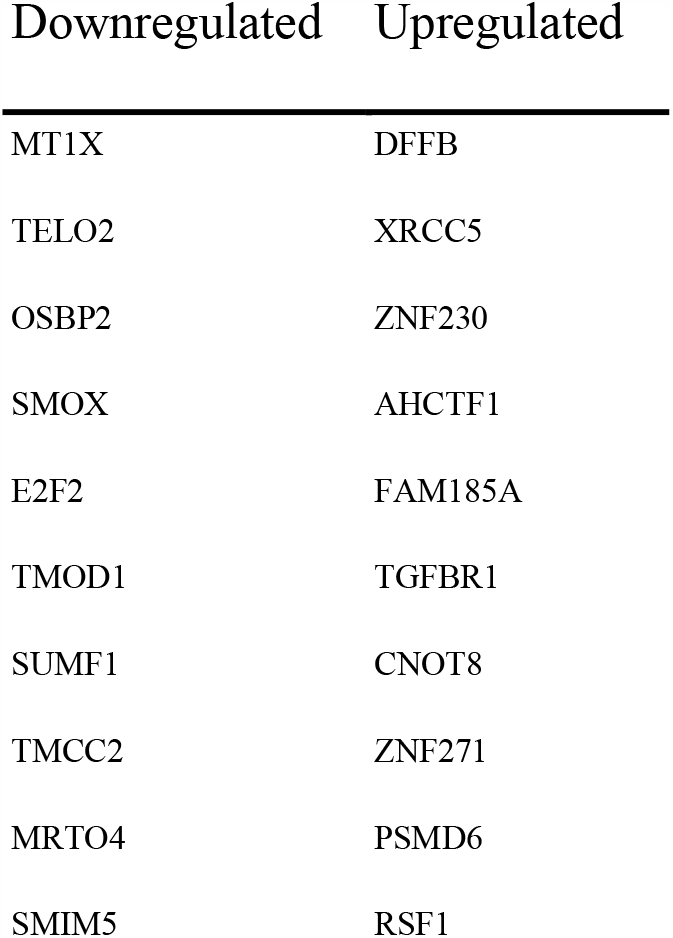
GSEA Ranked Genes List. (A) Top 10 downregulated and upregulated genes form GSEA ranked genes list

**Supplementary Table 2.** 250 gene Random Forest Classifier (A: Top) (B: Bottom)

|  | Precision | Recall | F1-score | Support |
| --- | --- | --- | --- | --- |
| AD | 0.969925 | 0.992308 | 0.980989 | 260 |
| CN | 0.388489 | 0.415385 | 0.401487 | 260 |
| EMCI | 0.518519 | 0.484615 | 0.500994 | 260 |
| LMCI | 0.509881 | 0.496154 | 0.502924 | 260 |
| Macro avg | 0.596703 | 0.597115 | 0.596598 | 1040 |
| Weighted avg | 0.596703 | 0.597115 | 0.596598 | 1040 |
| Accuracy | NA | NA | 0.597115 | 1040 |

Supplementary Table 2. 250 gene Random Forest Classifier (A: Top) (B: Bottom)
|  | Precision | Recall | F1-score | Support |
| --- | --- | --- | --- | --- |
| AD | 0.969925 | 0.992308 | 0.980989 | 260 |
| Control | 0.997416 | 0.989744 | 0.993565 | 780 |
| Macro avg | 0.983670 | 0.991026 | 0.987277 | 1040 |
| Weighted avg | 0.990543 | 0.990385 | 0.990421 | 1040 |
| Accuracy | NA | NA | 0.990385 | 1040 |

**Supplementary Table 3.**
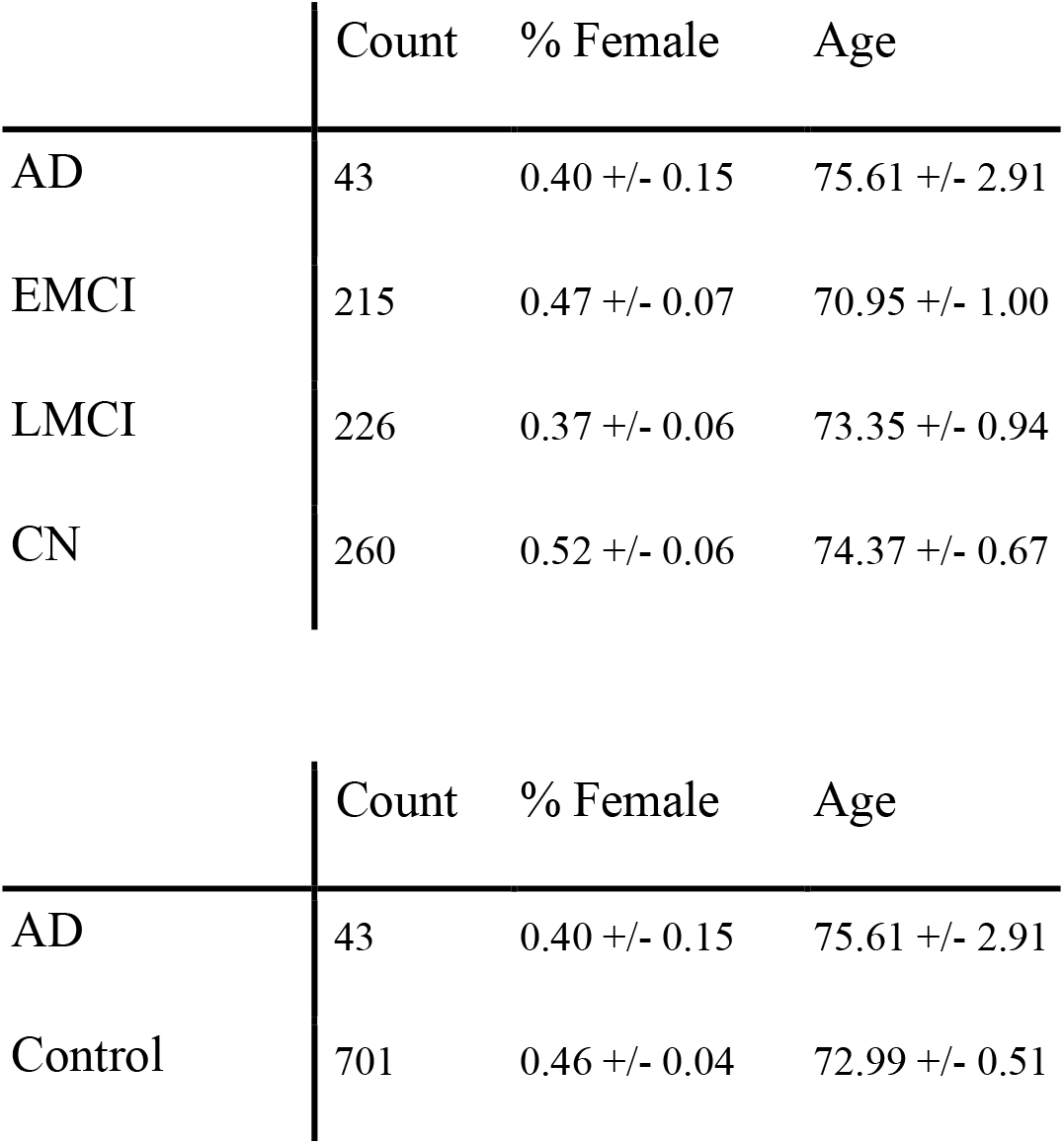
ADNI Demographics (A: Top) (B: Bottom) (A)Demographics of all 4 groups: CN, EMCI, LMCI, and AD (B)Binary Demographics: AD and Control

|  | Count | % Female | Age |
| --- | --- | --- | --- |
| AD | 43 | 0.40 +/- 0.15 | 75.61 +/- 2.91 |
| EMCI | 215 | 0.47 +/- 0.07 | 70.95 +/- 1.00 |
| LMCI | 226 | 0.37 +/- 0.06 | 73.35 +/- 0.94 |
| CN | 260 | 0.52 +/- 0.06 | 74.37 +/- 0.67 |
| AD | 43 | 0.40 +/- 0.15 | 75.61 +/- 2.91 |
| Control | 701 | 0.46 +/- 0.04 | 72.99 +/- 0.51 |

## 12. SUPPLEMENTARY FIGURES

**Supplementary Figure 1.**
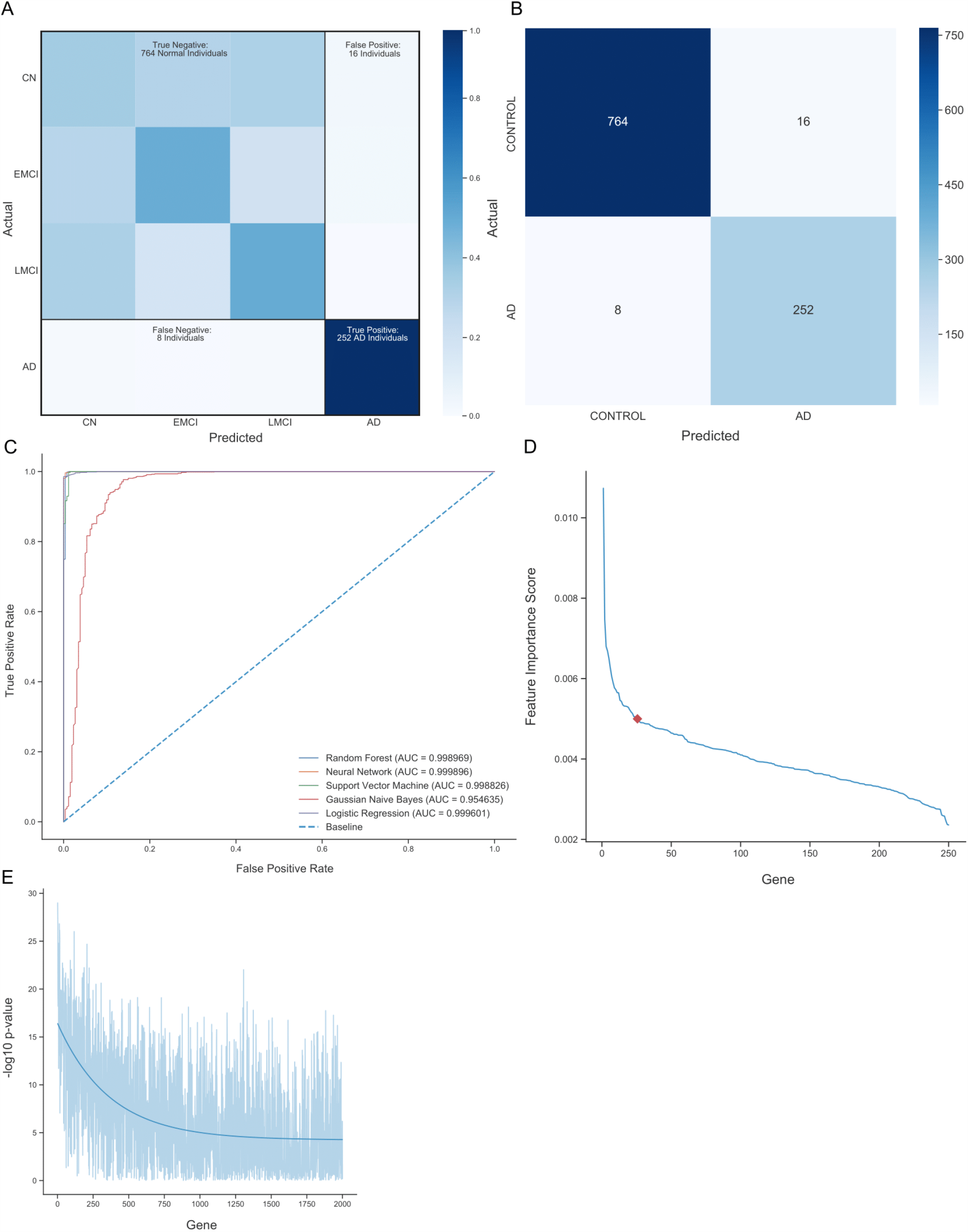
Random Forest Metrics. (A)Heatmap depicts results of a 5-fold cross-validation of the 25 gene RF classifier comprising 4 classes. Individuals that fall on the diagonal are classified according to their annotation (57.79% of individuals). Individuals that do not fall on the diagonal are mis-classified as a different diagnosis. (B)Confusion Matrix depicts results of a 5-fold cross-validation of the 25 gene RF classifier comparing AD to Control (CN, EMCI, LMCI). The sensitivity of detecting AD (true positive rate) is 96.9%. The specificity of detecting AD (true negative rate) is 97.95%. This data supports the high accuracy of the machine learning classifier. (C)Receiver operating characteristic curve for RF, Neural Network, Support Vector Machine, Gaussian Naive Bayes, and Logistic Regression on the top 250 gene’s in feature importance. (D)Line plot shows feature importance score of genes in the 250 gene RF classifier. Most of the feature importance is captured by the first 25 genes. (E)Line plot shows comparison of gene rank (according to feature importance) and -log10(Wilcox p-value). The trendline shows in general the higher the feature importance a gene has the lower its p-value.

**Supplementary Figure 2.**
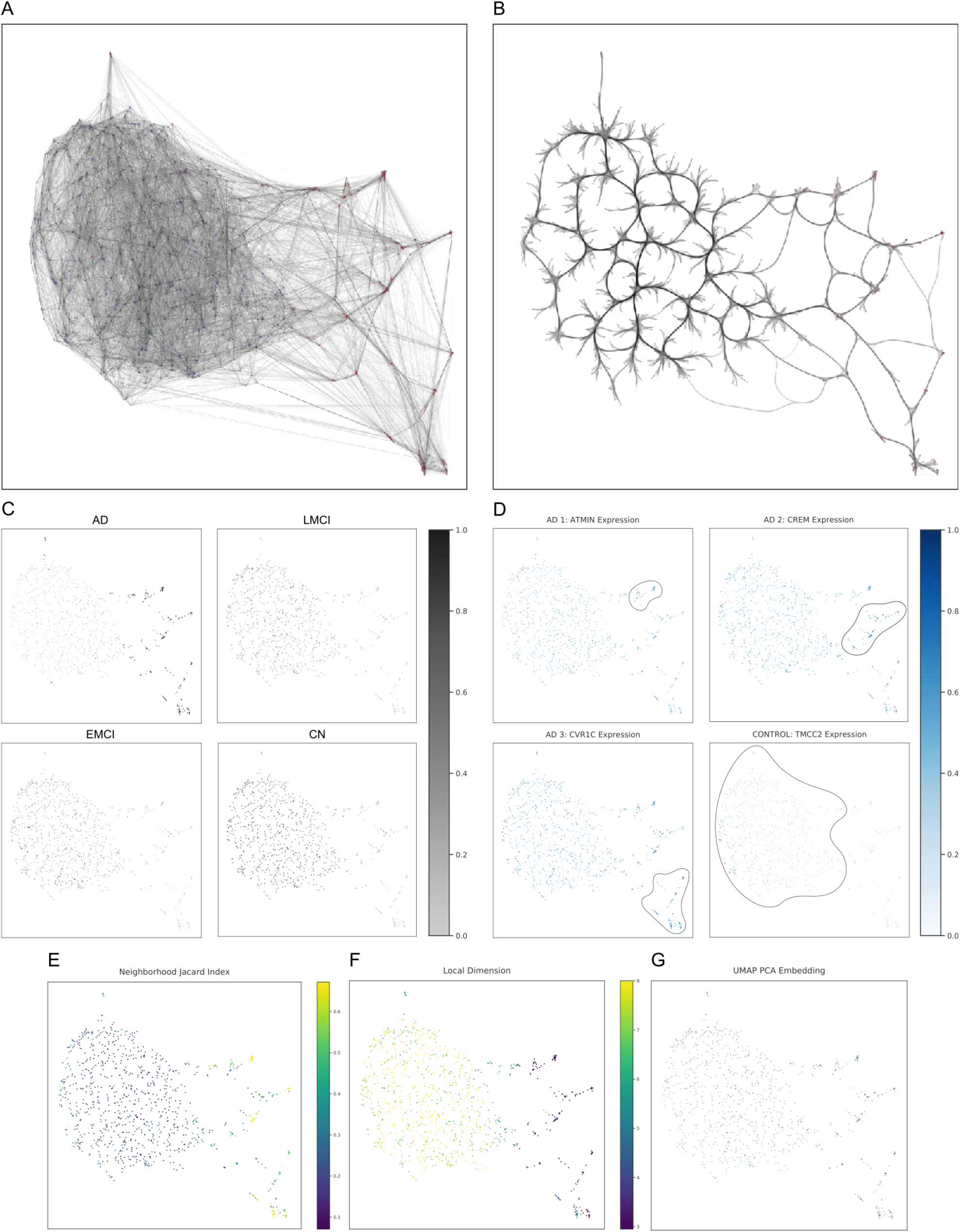
UMAP Metrics. (A)Connectivity visualization of UMAP (as in Figure 4D) shows connections between individuals. The UMAP structure was simplified down to a weighted graph and the connectivity in the manifold is visualized with respect to the embedding. (B)Edge-bundled connectivity visualization of UMAP (as in Figure 4D) shows main connections between individuals. Edge-bundling curves edges and groups nearby curves together to help convey structure. (C)UMAP visualization (as in Figure 4D) is overlaid with RF prediction scores. The further away an individual was from the main cluster the more confident RF was in its prediction of AD. These plots show that UMAP aligns well with the RF predictions. (D)UMAP visualization (as in Figure 4D) is overlaid with the relative expression levels of ATMIN, CREM, CVR1C, and TMCC2. These genes were identified as marker genes for the clusters AD1, AD2, AD3, and Control respectively using genes with the lowest p-value from wilcoxon rank sums test. These plots exemplify specific marker genes different among the three AD populations. (E-G) UMAP visualization (as in Figure 4D) is overlaid with diagnostics (Neighborhood Jaccard Index, Local Dimension, and PCA). For PCA, the first 3 PCs were converted into RGB values and overlaid. These plots confirm that UMAP maintains the global structure of the data.

**Supplementary Figure 3.**
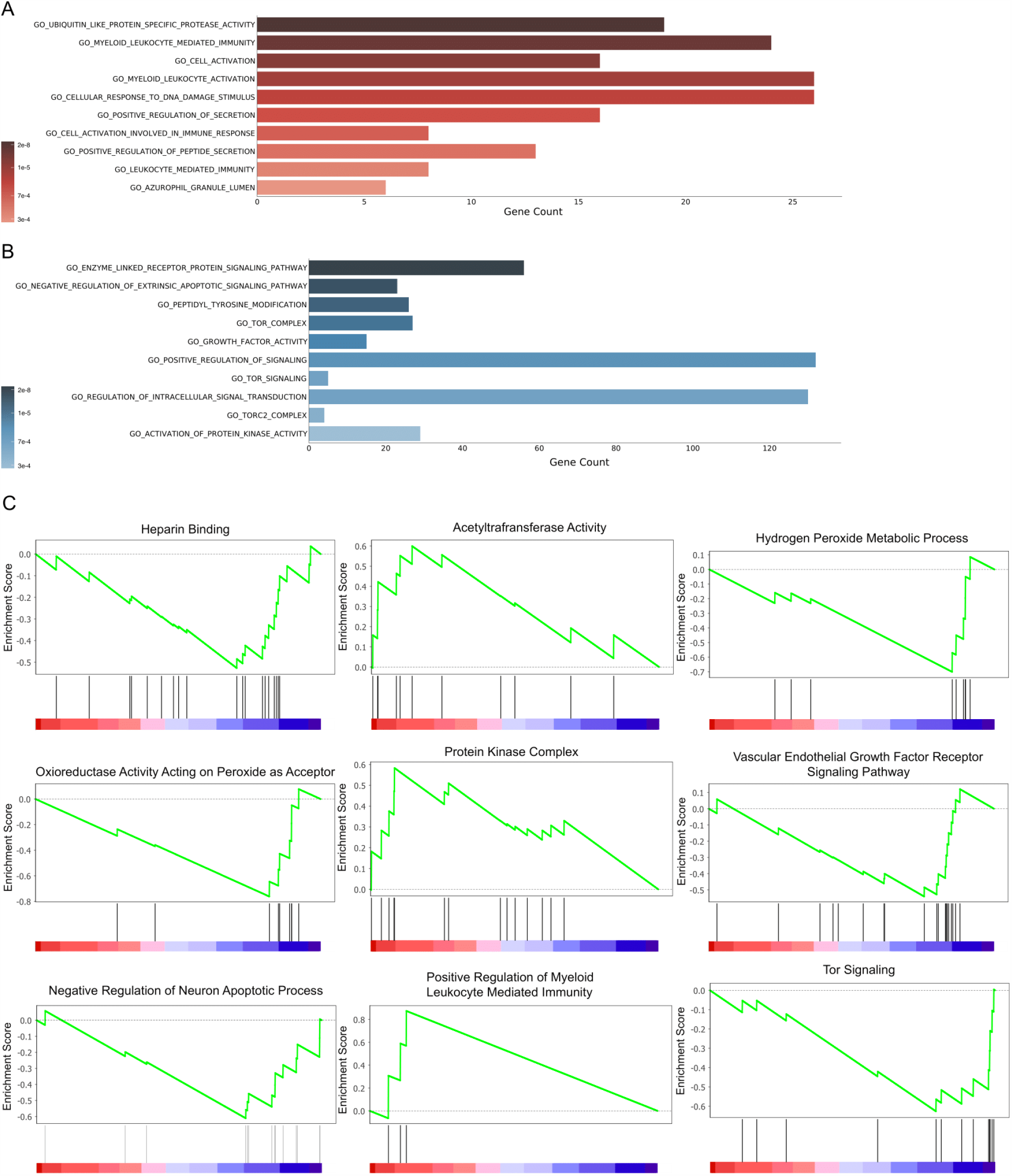
Enriched GO Terms. (A)Bar graph depicts the top 10 highly enriched pathways in AD according to the adjusted q-value. Length of the bar represents the gene set size while the color represents its q-value. (B)Bar graph depicts the top 10 highly enriched pathways in CN according to the adjusted q-value. Length of the bar represents the gene set size while the color represents its q-value. (C)Enrichment plots for 6 pathways downregulated (Left, Right) and 3 Pathways upregulated (Center) in AD. The horizontal axis are genes in order of their position in the ranked gene list and vertical lines represent gene hits.

## REFERENCES

[1] Burns A, Iliffe S. Alzheimer’s disease. BMJ. 2009;338:b158.

[2] Lozano R, Naghavi M, Foreman K, Lim S, Shibuya K, Aboyans V, Abraham J, Adair T, Aggarwal R, Ahn SY, et al. Global and regional mortality from 235 causes of death for 20 age groups in 1990 and 2010: a systematic analysis for the Global Burden of Disease Study 2010. Lancet. 2012;380(9859):2095–128

[3] Reitz C, Mayeux R. Alzheimer disease: epidemiology, diagnostic criteria, risk factors and biomarkers. Biochem Pharmacol. 2014;88(4):640–51

[4] Holtzman DM, Morris JC, Goate AM. Alzheimer’s disease: the challenge of the second century. Sci Transl Med. 2011;3(77):77sr1. doi:10.1126/scitranslmed.3002369

[5] Brion J, -P: Neurofibrillary Tangles and Alzheimer’s Disease. Eur Neurol 1998;40:130–140. doi: 10.1159/000007969

[6] Iqbal K, Liu F, Gong CX, Grundke-Iqbal I. Tau in Alzheimer disease and related tauopathies. Curr Alzheimer Res. 2010;7(8):656–664. doi:10.2174/156720510793611592

[7] Yashin AI, Fang F, Kovtun M, Wu D, Duan M, Arbeev K, et al. Hidden heterogeneity in Alzheimer’s disease: insights from genetic association studies and other analyses. Exp Gerontol. 2018;107:148–60.

[8] Evans, J. Grimley. Oxford Textbook of Geriatric Medicine. Oxford University Press, 2003.

[9] Shen L, Jia J. An Overview of Genome-Wide Association Studies in Alzheimer’s Disease. Neurosci Bull. 2016;32(2):183–190. doi:10.1007/s12264-016-0011-3

[10] Marian AJ. Molecular genetic studies of complex phenotypes. Transl Res. 2012;159:64–79. doi: 10.1016/j.trsl.2011.08.001.

[11] Raghavan N, Tosto G. Genetics of Alzheimer’s Disease: the Importance of Polygenic and Epistatic Components. Curr Neurol Neurosci Rep. 2017;17(10):78. Published 2017 Aug 21. doi:10.1007/s11910-017-0787-1

[12] Huynh RA, Mohan C. Alzheimer’s Disease: Biomarkers in the Genome, Blood, and Cerebrospinal Fluid. Front Neurol. 2017;8:102. Published 2017 Mar 20. doi:10.3389/fneur.2017.00102

[13] Small GW. Early diagnosis of Alzheimer’s disease: update on combining genetic and brain-imaging measures. Dialogues Clin Neurosci. 2000;2(3):241–246.

[14] Saykin AJ, Shen L, Foroud TM, et al. Alzheimer’s Disease Neuroimaging Initiative biomarkers as quantitative phenotypes: Genetics core aims, progress, and plans. Alzheimers Dement. 2010;6(3):265–273. doi:10.1016/j.jalz.2010.03.013

[15] Maes OC, Xu S, Yu B, Chertkow HM, Wang E, Schipper HM. Transcriptional profiling of Alzheimer blood mononuclear cells by microarray. Neurobiol Aging. 2007;28:1795–809.

[16] Liew CC, Ma J, Tang HC, Zheng R, Dempsey AA. The peripheral blood transcriptome dynamically reflects system wide biology: a potential diagnostic tool. J Lab Clin Med. 2006;147:126–32.

[17] Cosentino M, Colombo C, Mauri M, Ferrari M, Corbetta S, Marino F, et al. Expression of apoptosis-related proteins and of mRNA for dopaminergic receptors in peripheral blood mononuclear cells from patients with Alzheimer disease. Alzheimer Dis Assoc Disord. 2009;23:88–90.

[18] Saykin AJ, Shen L, Foroud TM, et al. Alzheimer’s Disease Neuroimaging Initiative biomarkers as quantitative phenotypes: Genetics core aims, progress, and plans. Alzheimers Dement. 2010;6(3):265–273. doi:10.1016/j.jalz.2010.03.013

[19] Shaw LM, Vanderstichele H, Knapik-Czajka M, et al. Cerebrospinal fluid biomarker signature in Alzheimer’s disease neuroimaging initiative subjects. Ann Neurol. 2009;65(4):403–413. doi:10.1002/ana.21610

[20] Somasundaram, Akila & Reddy, U. Srinivasulu. (2016). Data Imbalance: Effects and Solutions for Classification of Large and Highly Imbalanced Data.

[21] Chawla, N. V., et al. “SMOTE: Synthetic Minority Over-Sampling Technique.” Journal of Artificial Intelligence Research, vol. 16, 2002, pp. 321–357., doi:10.1613/jair.953.

[22] Kobak, Dmitry & Linderman, George. (2019). UMAP does not preserve global structure any better than t-SNE when using the same initialization. 10.1101/2019.12.19.877522.

[23] “About Tetrad.” Tetrad, www.phil.cmu.edu/tetrad/about.html.

[24] >“Greedy Fast Causal Inference (GFCI) Algorithm for Discrete Variables.” Center for Causal Discovery, www.ccd.pitt.edu//pdfs/GFCId.pdf.

[25] Hofer-Szabó, G. Relating Bell’s Local Causality to the Causal Markov Condition. Found Phys 45, 1110–1136 (2015). https://doi.org/10.1007/s10701-015-9868-7

[26] Weinberger, N. Faithfulness, Coordination and Causal Coincidences. Erkenn 83, 113–133 (2018). https://doi.org/10.1007/s10670-017-9882-6

[27] Jabbari F, Visweswaran S, Cooper GF. Instance-Specific Bayesian Network Structure Learning. Proc Mach Learn Res. 2018;72:169–180.

[28] Schmidt, Christopher, et al. “Order-Independent Constraint-Based Causal Structure Learning for Gaussian Distribution Models Using GPUs.” Proceedings of the 30th International Conference on Scientific and Statistical Database Management - SSDBM ‘18, 2018, doi:10.1145/3221269.3221292.

[29] Pearl, Judea, et al. Causal Inference in Statistics a Primer. Wiley, 2016.

[30] Ogarrio JM, Spirtes P, Ramsey J. A Hybrid Causal Search Algorithm for Latent Variable Models. JMLR Workshop Conf Proc. 2016;52:368–379.

[31] Database, Gene. “SUMF1 Gene (Protein Coding).” GeneCards, www.genecards.org/cgi-bin/carddisp.pl?gene=SUMF1&keywords=sumf1.

[32] Database, Gene. “KDM4B Gene (Protein Coding).” GeneCards, www.genecards.org/cgi-bin/carddisp.pl?gene=KDM4B.

[33] Database, Gene. “SMOX Gene (Protein Coding).” GeneCards, www.genecards.org/cgi-bin/carddisp.pl?gene=SMOX.

[34] “HLA-DPB1 Gene - Genetics Home Reference - NIH.” U.S. National Library of Medicine, National Institutes of Health, ghr.nlm.nih.gov/gene/HLA-DPB1.

[35] Wu, Cheng-Chia & Gupta, Tanush & Garcia, Victor & Ding, Yan & Schwartzman, Michal. (2013). 20-HETE and Blood Pressure Regulation Clinical Implications. Cardiology in review. 22. 10.1097/CRD.0b013e3182961659.

[36] National Center for Biotechnology Information. PubChem Database. NCBI Gene=1579, https://pubchem.ncbi.nlm.nih.gov/gene/CYP4A11/human (accessed on Feb. 25, 2020)

[37] “Solute Carrier Family 6 (Neurotransmitter Transporter, Gaba), Member 13; SLC6A13.” Error 403, www.omim.org/entry/615097.

[38] Database, Gene. “SLC6A13 Gene (Protein Coding).” GeneCards, www.genecards.org/cgi-bin/carddisp.pl?gene=SLC6A13.

[39] Database, Gene. “DYRK3 Gene (Protein Coding).” GeneCards, www.genecards.org/cgi-bin/carddisp.pl?gene=DYRK3.

[40] Stotani, S., Giordanetto, F., & Medda, F. (2016). DYRK1A inhibition as potential treatment for Alzheimer’s disease. Future Medicinal Chemistry, 8(6), 681–696. doi:10.4155/fmc-2016-0013

[41] Database, Gene. “LDB3 Gene (Protein Coding).” GeneCards, www.genecards.org/cgi-bin/carddisp.pl?gene=LDB3.

[42] “LDB3 Gene - Genetics Home Reference - NIH.” U.S. National Library of Medicine, National Institutes of Health, ghr.nlm.nih.gov/gene/LDB3.

[43] Young, Leah C., et al. “Kdm4b Histone Demethylase Is a DNA Damage Response Protein and Confers a Survival Advantage Following γ-Irradiation.” Journal of Biological Chemistry, vol. 288, no. 29, 2013, pp. 21376–21388., doi:10.1074/jbc.m113.491514.

[44] Database, Gene. “SUMF1 Gene (Protein Coding).” GeneCards, www.genecards.org/cgi-bin/carddisp.pl?gene=SUMF1.

[45] Malta, C. Di, et al. “Astrocyte Dysfunction Triggers Neurodegeneration in a Lysosomal Storage Disorder.” Proceedings of the National Academy of Sciences, vol. 109, no. 35, 2012, doi:10.1073/pnas.1209577109.

[46] Farkas E, Luiten PG. Cerebral microvascular pathology in aging and Alzheimer’s disease. Prog Neurobiol. 2001;64(6):575–611.

[47] Chen, Hao, et al. “A Machine Learning Method for Identifying Critical Interactions Between Gene Pairs in Alzheimer’s Disease Prediction.” Frontiers in Neurology, vol. 10, 2019, doi:10.3389/fneur.2019.01162.

[48] “Genetics.” Alzheimer’s Disease and Dementia, www.alz.org/alzheimers-dementia/what-is-alzheimers/causes-and-risk-factors/genetics.

[49] Park SY, Seo J, Chun YS. Targeted Downregulation of kdm4a Ameliorates Tau-engendered Defects in Drosophila melanogaster. J Korean Med Sci. 2019 Aug;34(33):e225. https://doi.org/10.3346/jkms.2019.34.e225.

[50] Aliseychik, M.P., Andreeva, T.V. & Rogaev, E.I. Immunogenetic Factors of Neurodegenerative Diseases: The Role of HLA Class II. Biochemistry Moscow 83, 1104–1116 (2018). https://doi.org/10.1134/S0006297918090122

[51] Price JL, Morris JC. Tangles and plaques in nondemented aging and preclinical Alzheimer disease. Ann Neurol 1999; 45:358–68.

[52] Bateman RJ, Xiong C, Benzinger TL, Fagan AM, Goate A, Fox NC, Marcus DS, Cairns NJ, Xie X, Blazey TM, Holtzman DM, Santacruz A, Buckles V, Oliver A, Moulder K, Aisen PS, Ghetti B, Klunk WE, McDade E, Martins RN, Masters CL, Mayeux R, Ringman JM, Rossor MN, Schofield PR, Sperling RA, Salloway S, Morris JC; the Dominantly Inherited Alzheimer Network. Clinical and Biomarker Changes in Dominantly Inherited Alzheimer’s Disease. N Engl J Med. 2012 Jul 11.

[53] Xu M, Zhang DF, Luo R, et al. A systematic integrated analysis of brain expression profiles reveals YAP1 and other prioritized hub genes as important upstream regulators in Alzheimer’s disease. Alzheimers Dement. 2018;14(2):215–229. doi:10.1016/j.jalz.2017.08.012

[54] Lee, T., Lee, H. Prediction of Alzheimer’s disease using blood gene expression data. Sci Rep 10, 3485 (2020). https://doi.org/10.1038/s41598-020-60595-1

[55] Benjamini Y, Hochberg Y. Controlling the false discovery rate: a practical and powerful approach to multiple testing. Journal of the Royal Statistical Society B. 1995;57:289–300.

[56] Shannon P, Markiel A, Ozier O, et al. Cytoscape: a software environment for integrated models of biomolecular interaction networks. Genome Res. 2003;13(11):2498–2504. doi:10.1101/gr.1239303

[57] Kucera M, Isserlin R, Arkhangorodsky A, Bader GD. AutoAnnotate: A Cytoscape app for summarizing networks with semantic annotations. F1000Res. 2016;5:1717. Published 2016 Jul 15. doi:10.12688/f1000research.9090.1

[58] Subramanian A., Tamayo P., Mootha V.K., Mukherjee S., Ebert B.L., Gillette M.A., Paulovich A., Pomeroy S.L., Golub T.R., Lander E.S., Mesirov J.P. Gene set enrichment analysis: a knowledge-based approach for interpreting genome-wide expression profiles. Proc. Natl. Acad. Sci. 2005;102(43):15545–15550.

[59] Database, Gene. “SMOX Gene (Protein Coding).” GeneCards, https://www.genecards.org/cgi-bin/carddisp.pl?gene=HLA-DPB1

[60] Monsonego A, Nemirovsky A, Harpaz I. CD4 T cells in immunity and immunotherapy of Alzheimer’s disease. Immunology. 2013;139(4):438–446. doi:10.1111/imm.12103

[61] Ariga T, Miyatake T, Yu RK. Role of proteoglycans and glycosaminoglycans in the pathogenesis of Alzheimer’s disease and related disorders: amyloidogenesis and therapeutic strategies--a review. J Neurosci Res. 2010;88(11):2303–2315. doi:10.1002/jnr.22393

[62] Govindpani K, McNamara LG, Smith NR, et al. Vascular Dysfunction in Alzheimer’s Disease: A Prelude to the Pathological Process or a Consequence of It?. J Clin Med. 2019;8(5):651. Published 2019 May 10. doi:10.3390/jcm8050651

[63] Shekhar, S., Varghese, K., Li, M., Fan, L., Booz, G. W., Roman, R. J., & Fan, F. (2019). Conflicting roles of 20-HETE in hypertension and stroke. International journal of molecular sciences, 20(18), [4500]. https://doi.org/10.3390/ijms20184500

[64] Magi S, Castaldo P, ML Macrì, Maiolino M, Matteucci A, Bastioli G, Gratteri S, Amoroso S, Lariccia V. Intracellular Calcium Dysregulation: Implications for Alzheimer’s Disease. Biomed Res Int. 2016;2016 6701324. doi:10.1155/2016/6701324. PMID: 27340665; PMCID: PMC4909906.

[65] Di Wu, Kousuke Noda, Miyuki Murata, Ye Liu, Atsuhiro Kanda, Susumu Ishida; Modulation of Spermine Oxidation in Müller Glial Cells under Hypoxic Condition. Invest. Ophthalmol. Vis. Sci. 2017;58(8):5199.

[66] Christen Y Am J Clin Nutr. 2000 Feb; 71(2):621S–629S.

[67] Huang WJ, Zhang X, Chen WW. Role of oxidative stress in Alzheimer’s disease. Biomed Rep. 2016;4(5):519–522. doi:10.3892/br.2016.630

[68] Amyloid diseases: abnormal protein aggregation in neurodegeneration. Koo EH, Lansbury PT Jr, Kelly JW Proc Natl Acad Sci U S A. 1999 Aug 31; 96(18):9989–90.

[69] Grundke-Iqbal I, Iqbal K, Quinlan M, Tung YC, Zaidi MS, Wisniewski HM. Microtubule-associated protein tau. A component of Alzheimer paired helical filaments. J Biol Chem. 1986;261(13):6084–9.

[70] Wischik CM, Novak M, Edwards PC, Klug A, Tichelaar W, Crowther RA. Structural characterization of the core of the paired helical filament of Alzheimer disease. Proc Natl Acad Sci U S A. 1988;85(13):4884–8.

[71] Database, Gene. “SMOX Gene (Protein Coding).” GeneCards, https://www.genecards.org/cgi-bin/carddisp.pl?gene=DFFB

[72] Kolobov, V.V., Davydova, T.V., Zakharova, I.A. et al.. Glutamate antibodies repress expression of Dffb gene in brain of rats in experimental Alzheimer’s disease. Mol Biol 46, 678–686 (2012). https://doi.org/10.1134/S0026893312040061

[73] Di Malta C, Fryer JD, Settembre C, Ballabio A. Autophagy in astrocytes: a novel culprit in lysosomal storage disorders. Autophagy. 2012;8(12):1871–1872. doi:10.4161/auto.22184

[74] Antonio Di Meco, Mary Elizabeth Curtis, Elisabetta Lauretti, Domenico Praticò, Autophagy Dysfunction in Alzheimer’s Disease: Mechanistic Insights and New Therapeutic Opportunities, Biological Psychiatry, Volume 87, Issue 9, 2020, Pages 797–807, ISSN 0006-3223, https://doi.org/10.1016/j.biopsych.2019.05.008.

[75] Calissano P, Matrone C, Amadoro G. Apoptosis and in vitro Alzheimer disease neuronal models. Commun Integr Biol. 2009;2(2):163–169. doi:10.4161/cib.7704

[76] McInnes, L, Healy, J, UMAP: Uniform Manifold Approximation and Projection for Dimension Reduction, ArXiv e-prints 1802.03426, 2018

[77] Ogarrio JM, Spirtes P, Ramsey J. A Hybrid Causal Search Algorithm for Latent Variable Models. JMLR Workshop Conf Proc. 2016;52:368–379.

[78] M.T. Ferretti, M. Merlini, C. Spni, C. Gericke, N. Schweizer, G. Enzmann, B. Engelhardt, L. Kulic, T. Suter, R.M. Nitsch, T-cell brain infiltration and immature antigen-presenting cells in transgenic models of Alzheimer’s disease-like cerebral amyloidosis, Brain, Behavior, and Immunity, Volume 54, 2016, Pages 211–225, ISSN 0889-1591, https://doi.org/10.1016/j.bbi.2016.02.009.

[79] Gary Beecham, Kara Hamilton, Gerard Schellenberg, Margaret Pericak-Vance, Thomas Montine Neurology Apr 2014, 82 (10 Supplement) S28.002;

[80] Moradifard, Shirin, et al. “Analysis of MicroRNA and Gene Expression Profiles in Alzheimer’s Disease: A Meta-Analysis Approach.” Scientific Reports, vol. 8, no. 1, 2018, doi:10.1038/s41598-018-20959-0.

